# Triplex and other DNA motifs show motif-specific associations with mitochondrial DNA deletions and species lifespan

**DOI:** 10.1101/2020.11.13.381475

**Authors:** Kamil Pabis

## Abstract

The “theory of resistant biomolecules” posits that long-lived species show resistance to molecular damage at the level of their biomolecules. Here, we test this hypothesis in the context of mitochondrial DNA (mtDNA) as it implies that predicted mutagenic DNA motifs should be inversely correlated with species maximum lifespan (MLS).

First, we confirmed that guanine-quadruplex and direct repeat (DR) motifs are mutagenic, as they associate with mtDNA deletions in the human major arc of mtDNA, while also adding mirror repeat (MR) and intramolecular triplex motifs to a growing list of potentially mutagenic features. What is more, triplex motifs showed disease-specific associations with deletions and an apparent interaction with guanine-quadruplex motifs.

Surprisingly, even though DR, MR and guanine-quadruplex motifs were associated with mtDNA deletions, their correlation with MLS was explained by the biased base composition of mtDNA. Only triplex motifs negatively correlated with MLS even after adjusting for body mass, phylogeny, mtDNA base composition and effective number of codons.

Taken together, our work highlights the importance of base composition for the comparative biogerontology of mtDNA and suggests that future research on mitochondrial triplex motifs is warranted.

## INTRODUCTION

Macromolecular damage to lipids, proteins and DNA accumulates with aging (**Richardson and Schadt 2014, Gladyshev 2013**), whereas cells isolated from long-lived species are resistant to genotoxic and cytotoxic drugs, giving rise to the multistress resistance theory of aging (**Miller 2009, Hamilton and Miller 2016**). By extension of this idea, the “theory of resistant biomolecules” posits that lipids, proteins and DNA itself should be resilient in long-lived species (**Pamplona and Barja 2007**). In support of this theory, it was shown that long-lived species possess membranes that contain fewer lipids with reactive double bonds (**Valencak and Ruf 2007**) and perhaps a lower content of oxidation-prone cysteine and methionine in mitochondrially encoded proteins (see **Aledo et al. 2012** for a discussion).

Mitochondrial DNA (mtDNA) mutations constitute one type of macromolecular damage that accumulates over time. Point mutations accumulate in proliferative tissues like the colon and in some progeroid mice (**Kauppila et al. 2017**), while the accumulation of mtDNA deletions in postmitotic tissues may underpin certain age-related diseases like Parkinson’s and sarcopenia (**Lawless et al. 2020, Bender et al. 2006**).

If the theory of resistant biomolecules can be generalized, the mtDNA of long-lived species should resist both point mutation and deletion formation. However, we will focus on deletions because they are more pathogenic than point mutations at the same level of heteroplasmy (**Gamamge et al. 2014**) and human tissues do not accumulate high levels of point mutations observed in progeroid mouse models (**Khrapko et al. 2006**).

Since deletion formation depends on the primary sequence of the mtDNA (sequence motifs) it is amenable to bioinformatic methods. Ever since a link between direct repeat (DR) motifs and deletion formation became known, variations of the theory of resistant biomolecules have been tested, although not necessarily under this name. It was reasoned that long-lived species evolved to resist deletion formation and mtDNA instability by reducing the number of mutagenic motifs in their mtDNA (**Khaidakov et al. 2006, Yang et al. 2013**).

We aim to extend these findings by re-evaluating and establishing new candidate motifs, which we then correlate with species maximum lifespan (MLS). Studying multiple motif classes at once also allows us to reveal relationships between potentially overlapping mtDNA motifs that may affect the data. We define candidate motifs as those that are associated with deletion formation inside the major arc of human mtDNA, because during asynchronous replication the major arc is single stranded for extended periods of time (**Persson et al. 2019**) which should favor the formation of secondary structures. Finally, we test if these motifs correlate with the MLS of mammals, birds and ray-finned fishes after correcting for potential biases, especially global mtDNA base composition which is an important confounder (**Aledo et al. 2012**) yet is neglected in some studies (**Yang et al. 2013**).

The choice of motifs to study is based on biological plausibility and published literature that will be briefly reviewed below. Mutagenic motifs include repeats as well as guanine-quadruplex (GQ)- and triplex-forming motifs. DR motifs can lead to DNA instability through strand-slippage if two DR motifs mispair during replication (**Persson et al. 2019**). Whereas inverted repeat (IR), G-quadruplex and triplex motifs destabilize progression of the replication fork through the formation of stable secondary structures. Some of the structures formed include hairpins for IR motifs (**Tremblay-Belzile et al. 2015**), triple stranded DNA for triplex motifs and bulky stacks of guanines for G-quadruplex motifs (**Bacolla et al. 2016; Fig. 1**). Mirror repeat (MR) and everted repeat (ER) motifs, in contrast, do not allow stable Watson-Crick base pairing and are thus less likely to be mutagenic, although a subset of MR motifs may form triplex structures (**Kamat et al. 2016)**.

**Figure 1.**
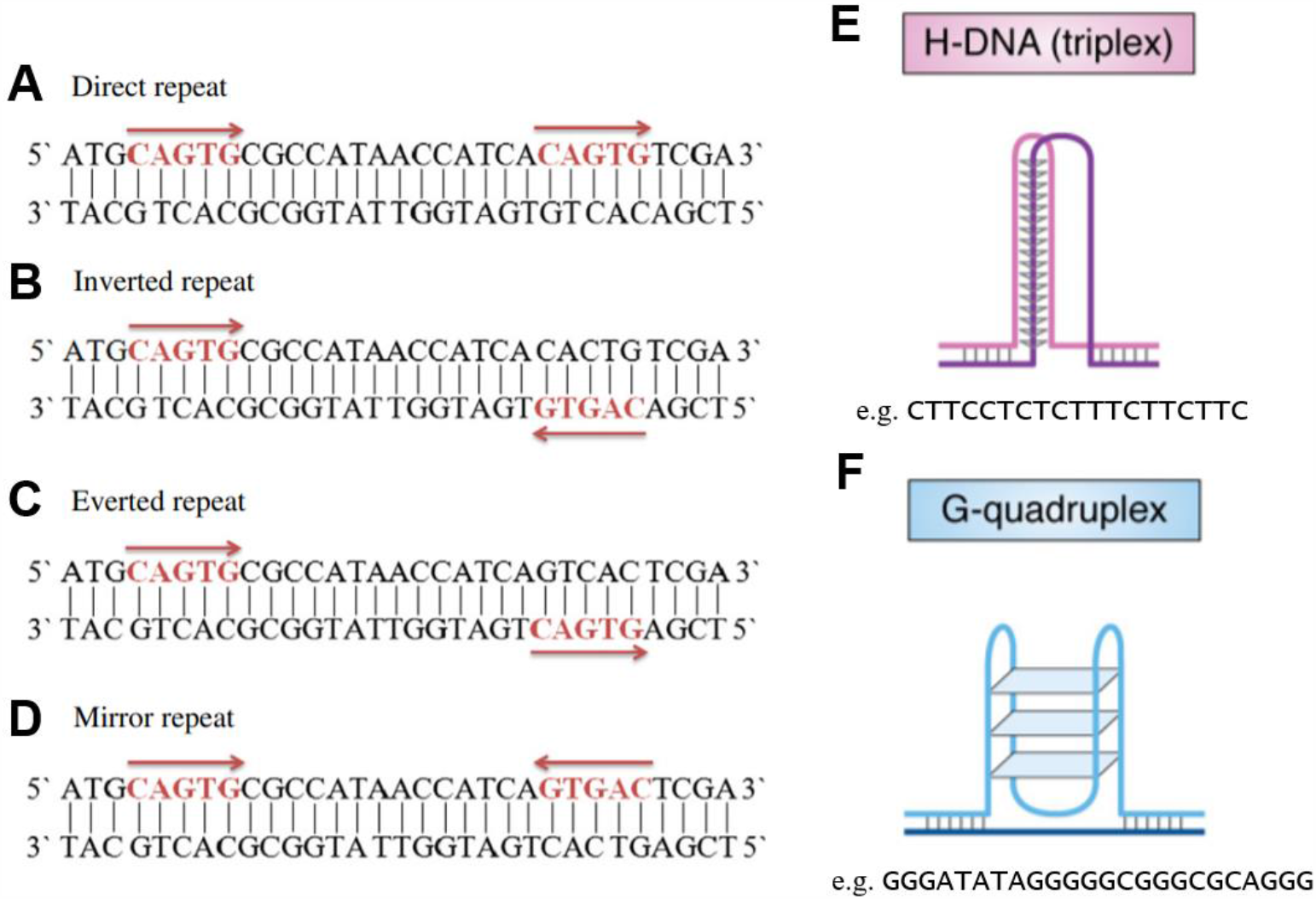
A. Direct repeat, both half-sites have the same orientation. B. Inverted repeat, the half-sites are complementary and has mirror symmetry. C. Everted repeat, the half-sites are complementary. D. Mirror repeat, the half-sites have mirror symmetry. E. Triplex motifs can form a triple helical DNA structure also called H-DNA. F. In a G-quadruplex multiple G-quartets (depicted as blue rectangles) stack on top of each other.

Thus, many motifs can be mutagenic in principle, but what is the evidence that these motifs are related to mtDNA instability, particularly deletions, and MLS?

Paradoxically, while DRs are the motif most consistently associated with mtDNA deletion breakpoints (BPs), despite preliminary reports (**Khaidakov et al. 2006, Lakshmanan et al. 2012, Yang et al. 2013**), no correlation with species MLS was seen in recent studies (**Lakshmanan et al. 2015**). In contrast, with the exception of one preprint (**Mikhailova et al. 2020**), IRs are not known to be associated with mtDNA deletions (**Dong et al. 2014**), although they do show a negative relationship with species MLS (**Yang et al. 2013**) and may contribute to inversions (**Tremblay-Belzile et al. 2015**). Whether age-related mtDNA inversions underlie any pathology, however, requires further study. Finally, G-quadruplex motifs are associated with both deletions (**Dong et al. 2014**) and point mutations (**Butler et al. 2020**), but no study tested if they correlate with MLS. Triplex motifs are poorly studied with one report finding no association between these motifs and deletions (**Oliveira et al. 2013**).

Based on these studies we decided to test the theory of resistant biomolecules by quantifying DR, MR, IR, ER, G-quadruplex- and triplex-forming motifs. We stipulate that if a motif class played a causal role in aging, it should be involved in deletion formation and its abundance should be negatively correlated with species MLS.

Adapted from **Gurusaran et al. (2013)** and **Khristich and Mirkin (2020**) with permission. Half-sites shown in red.

## METHODS

### Detection of DNA motifs

Repeats were detected by a script written in R (vR-3.6.3). Briefly, to find all repeats with N basepairs (bps), the mtDNA light strand is truncated by 0 to N bps and each of the N truncated mtDNAs is then split every N bps. This generates every possible substring (and thus repeat) of length N. In the next step, duplicate strings are removed. Afterwards we can find DR (a substring with at least two matches in the mtDNA), MR (at least one match in the mtDNA and on its reverse), IR (at least one match in the mtDNA and on its reverse-complement) and ER motifs (at least one match in the mtDNA and on its complement). Overlapping and duplicate repeats were not counted for the correlation between repeats and MLS. The code for the analyses performed in this paper can be found on github (pabisk/aging_triplex2).

Unless stated otherwise, all analyses were performed in R. G-quadruplex motifs were detected by the pqsfinder package (v2.2.0, **Hon et al. 2017**). Intramolecular triplex-forming motifs were detected by the triplex package (v1.26.0, **Hon et al. 2013**) and duplicates were removed. We also compared the data with two other publicly available tools, Triplexator (**Buske et al. 2013**), and with the non-B DNA motif search tool (nBMST; **Cer et al. 2011**). Triplexator was run on a virtual machine in an Oracle VM VirtualBox (v6.1) in -ss mode on the human mitochondrial genome and its reverse complement, the results were combined and overlapping motifs from the output were removed. We used the web interface of nBMST to detect mirror repeats/triplexes (v1.0).

### Association between motifs and major arc deletions

The major arc was defined as the region between position 5747 and 16500 of the human mtDNA (NC_012920.1). The following deletions and their breakpoints were located in this region and included: 1066 deletions from the MitoBreak database (**Damas et al. 2014, mtDNA Breakpoints**.**xlsx**), 1114 from **Persson et al. (2019)** and 1894 from **Hjelm et al. (2019)**.

Each deletion is defined by two breakpoints. A breakpoint pair was considered to associate with a motif if the motif fell within a defined window around one or both breakpoints, depending on the analysis. The window size was chosen in relation to the length of the studied motifs (30 bp for repeats and 50 bp for other motifs).

Three different motif orientations relative to the breakpoints were considered. Two orientations for motifs with half-sites (i.e. repeats), either both half-sites at any one breakpoint of a deletion, or one half-site per breakpoint of a deletion. Motifs with overlapping half-sites were not counted. In the third case, distinct G-quadruplex and triplex motifs could associate with one or both breakpoints of a deletion, but were at most counted once, since the latter case is sufficiently rare.

In order to exclude overlapping “hybrid” motifs, MR and DR motifs with the same sequence were removed whereas triplex and G-quadruplex motifs were removed if they were in proximity.

To generate controls, the mtDNA deletions as a whole were randomly redistributed inside the major arc which, because of the fixed deletion size, allowed us to approximate the original distribution of breakpoints (as suggested by **Oliveira et al. 2013**). Significance was determined via one-sample t-test in Prism (v7.04) by comparing actual breakpoints to 20 such randomized controls. Alternative controls were generated by shifting each breakpoint by 200 bp towards the midpoint of the major arc or as in **Fig. S9**.

### Cancer associated breakpoints

We obtained all autosomal breakpoints available from the Catalogue Of Somatic Mutations In Cancer (COSMIC; release v92, 27^th^ August 2020), which includes deletions, inversions, duplications and other abnormalities (n=587515 in total). After removing breakpoints whose sequences could not be retrieved (<1.7%), we quantified the number of predicted G-quadruplex and triplex motifs in a 500 bp window centered on the breakpoints using default settings for the detection of these motifs. Sequences of breakpoint regions were obtained from the GRCh38 build of the human genome using the BSgenome package (v1.3.1). Each breakpoint shifted by +3000 bps served as its own control.

### Lifespan, base composition and life history traits

We included three phylogenetic classes in our analysis for which we had sufficient data (n>100), mammals, birds and ray-finned fishes (actinopterygii). MLS and body mass were determined from the AnAge database (**Tacutu et al. 2018**) and, for mammals, supplemented with data from **Pacifici et al. (2013)**. The mtDNA accessions were obtained from an updated version of MitoAge (unpublished; **Toren et al. 2016**). Species were excluded if body mass data was unavailable, if the sequence could not be obtained using the genbankr package (v1.14.0), or if the extracted cytochrome B DNA sequence did not allow for an alignment, precluding phylogenetic correction. The species data can be found in the supplementary (**Species Data**.**xlsx**).

We analyzed the full mtDNA sequence, heuristically defined as the mtDNA sequence between the first and last encoded tRNA, excluding the D-loop, which is rarely involved in repeat-mediated deletion formation (**Yang et al. 2013**). The effective number of codons was calculated using Wright’s Nc (**Smith et al. 2019**). Base composition was calculated for the light-strand. GC skew was calculated as the fraction (G − C)/(G + C) and AT skew as (A − T)/(A + T). All correlations are Pearson’s R. Partial correlations were performed using the ppcor package (v1.1).

### Phylogenetic generalised least squares and phylogenetic correction

Observed correlations between traits and lifespan can be spurious due to shared species ancestry (**Speakman 2005**). To correct for this, we use phylogenetic generalised least squares (PGLS) implemented in the caper package (v1.0.1). Species phylogenetic trees were constructed via neighbor joining based on aligned cytochrome B DNA sequences using Clustal Omega from the msa package (v1.18.0) and in the resulting mammalian and bird tree, four branch edge lengths were equal to zero, which were set to the lowest non-zero value in the dataset.

## RESULTS

### Direct repeats and mirror repeats are over-represented at mtDNA deletion breakpoints

In order to define candidate mtDNA motifs that could be linked with lifespan, we started by reanalyzing motifs that associate with mtDNA deletion breakpoints reported in the MitoBreak database (**Damas et al. 2014; Fig. S1; mtDNA Breakpoints**.**xlsx**). In the below analysis, we consider DR and IR motifs thought to be mutagenic, as well as MR and ER motifs, so far not known to be mutagenic and we pool all 6 to 15 bp long repeats, since the data is similar between different repeat lengths (**Fig. S2**).

As shown by others, we found that DR motifs often flank mtDNA deletions (**Fig. 2A**). In contrast, no strong association was seen for ER and IR motifs, even considering a larger window around the breakpoint to allow for the fact that IRs could bridge and destabilize mtDNA over long distances (**Persson et al. 2019; Fig. S3**).

**Figure 2.**
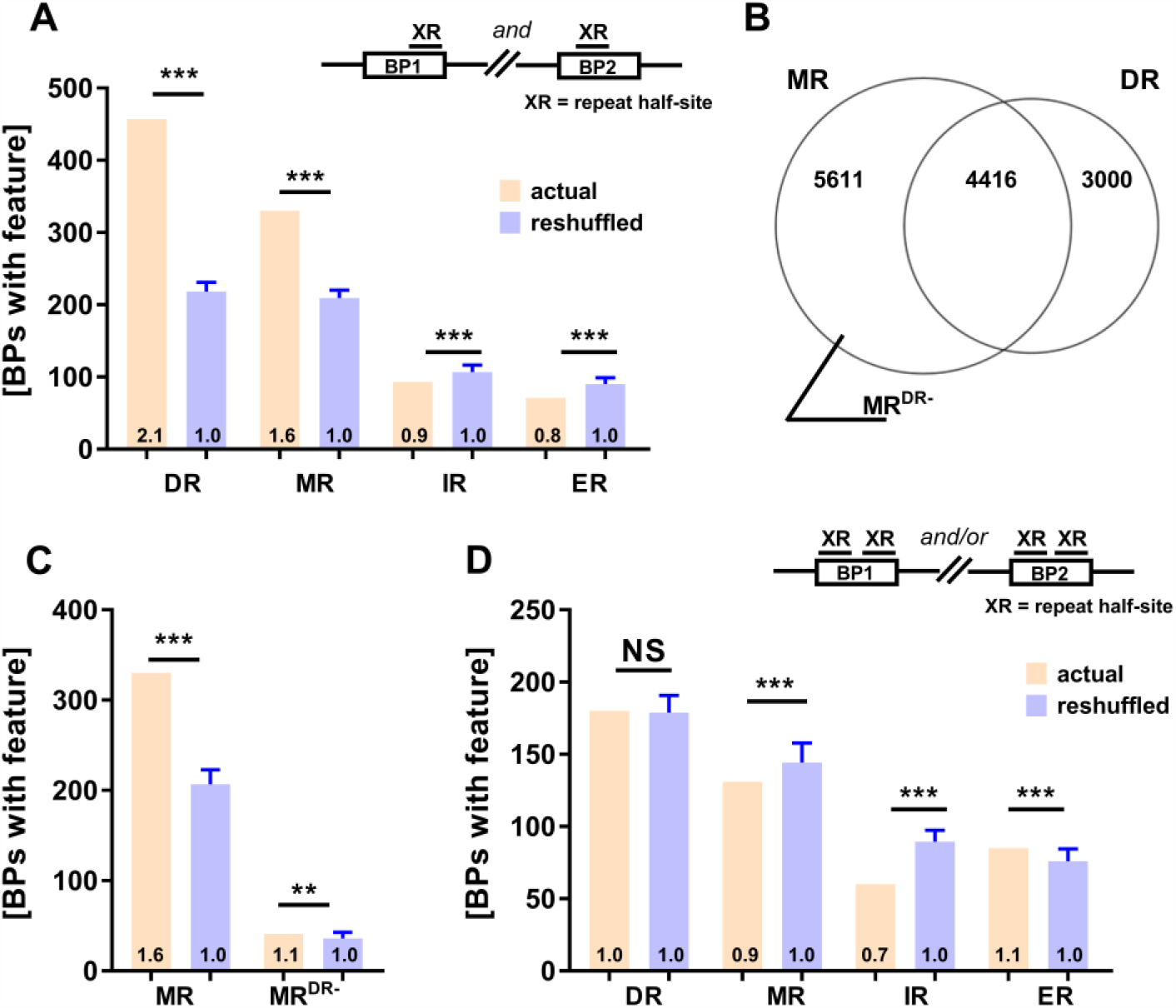
Direct repeat (DR) and mirror repeat (MR) motifs are significantly enriched around actual deletion breakpoints (BPs) compared to reshuffled BPs, but the same is not true for inverted repeat (IR) and everted repeat (ER) motifs (A, D). The surprising correlation between MR motifs and deletion BPs is attenuated when MRs that have the same sequence as DR motifs are removed (B, C). Controls were generated by reshuffling the deletion BPs while maintaining their distribution (n=20, mean ±SD shown). The schematic drawings above (A, D) depict the orientation of the repeat (XR) half-sites in relation to the BPs. *** p < 0.001; ** p < 0.01 by one sample t-test. A) The number of deletions associated with DR, MR, IR or ER motifs at both BPs compared with reshuffled controls. B) Venn diagram showing the number of MR, DR and hybrid MR-DR motifs that were identified within the major arc. C) The number of deletions associated with MR motifs, before (MR) and after removal of hybrid MR-DR motifs (MR^DR-^), compared with reshuffled controls. D) The number of deletions associated with DR, MR, IR or ER motifs at either BP compared with reshuffled controls.

Surprisingly, we also found MR motifs flanking deletion breakpoints more often than expected by chance (**Fig. 2A**). However, DR and MR motifs are known to correlate with each other (**Shamanskiy et al. 2019; Fig. 5B**) and indeed we noticed a large sequence overlap between MR and DR motifs (**Fig. 2B**), which could explain an apparent over-representation of MRs at breakpoints. Removal of overlapping MR-DR hybrid motifs confirmed this suspicion. After this correction, the degree of enrichment was strongly attenuated (**Fig. 2C**) and the total number of breakpoints flanked by MR motifs was reduced by >80%. Nevertheless, long MR motifs remained particularly over-represented around deletions (**Fig. S4**).

Since the prior analysis only considered motifs that flank both breakpoints, we next tested the idea that IR and other motifs could be mutagenic if both half-sites are found at any of the breakpoints. However, in this analysis no motif class showed enrichment around breakpoints (**Fig. 2D**).

### Predicted triplex-forming motifs are over-represented at mtDNA breakpoints

Given the association between MR motifs and breakpoints we decided to analyze triplex motifs, a special case of homopurine and homopyrimidine mirror repeats (**Khristich and Mirkin 2020, Bissler 2007**), and their association with deletion breakpoints in the MitoBreak database.

Here, we use the triplex package to predict intramolecular triplex motifs because it has several advantages compared to other software (**Hon et al. 2013**). For example, using the nBMST tool, as in a previous study of mtDNA instability (**Oliveira et al. 2013**), we only identified two potential triplex motifs within the major arc that did not overlap with the six motifs identified by the triplex package (**Table S1**). In contrast, using Triplexator (**Buske al. 2013**) we were able to detect four of the six triplex motifs and the motifs detected by Triplexator were also enriched at breakpoints (**Table S2**).

We noticed that predicted triplexes are G-rich and thus could be related to G-quadruplex motifs (**Doluca et al. 2013**). In a comparison of the two motif types, however, we found several differences (**Table S1, S3**). Triplex motifs were shorter and less abundant than predicted G-quadruplexes, associated with fewer breakpoints altogether (**Fig. 3**) and, in contrast to G-quadruplexes almost exclusive to the G-rich mtDNA heavy-strand, triplex motifs were also common on the light-strand.

**Figure 3.**
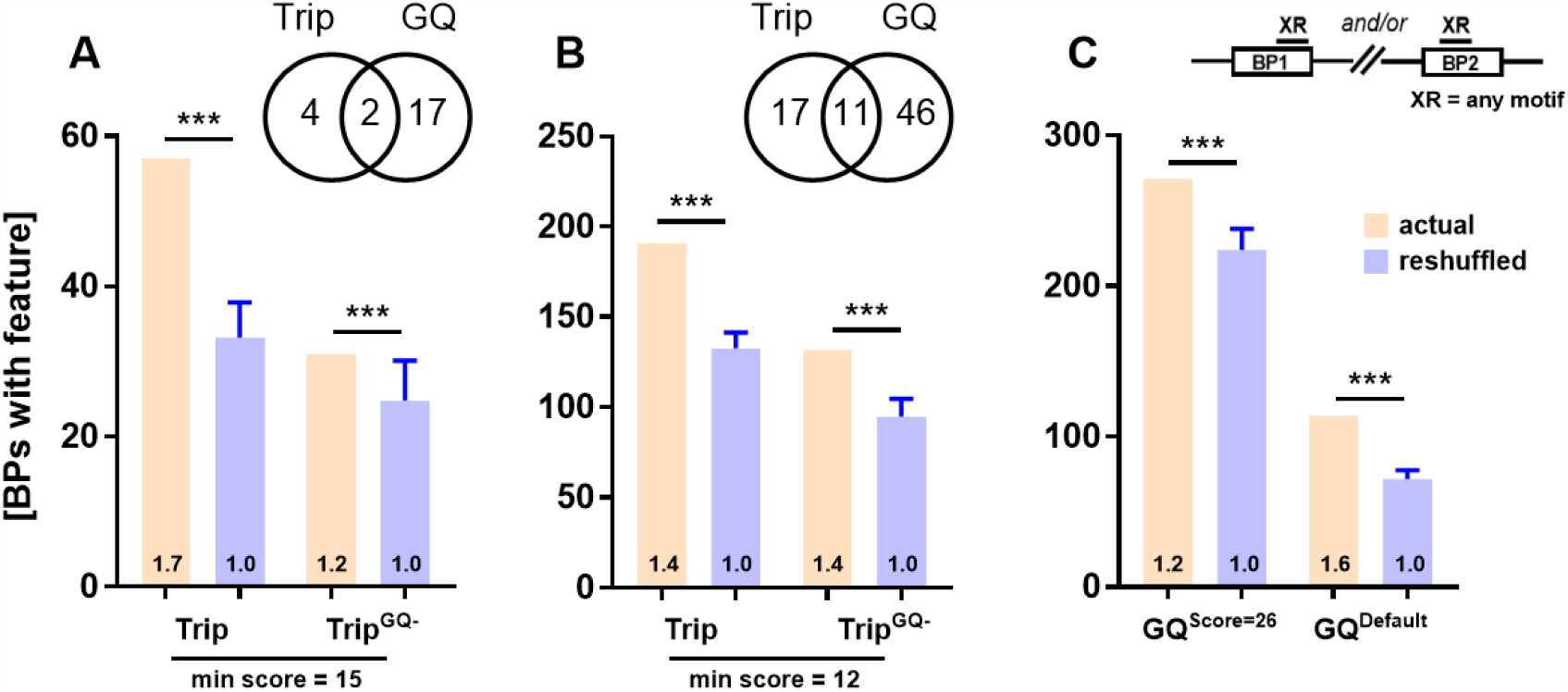
Triplex motifs are significantly enriched around actual breakpoints (BPs) compared to reshuffled BPs (A, B) even after removal of G-quadruplex (GQ)-triplex hybrid motifs (Trip^GQ-^). The number of unique triplex motifs, GQ motifs and of hybrid triplex-GQ motifs, within the mtDNA major arc, is shown in the Venn diagrams above (A, B). Enrichment of GQ motifs around BPs is shown for comparison in (C). Controls were generated by reshuffling the deletion BPs while maintaining their distribution (n=20, mean ±SD shown). The schematic drawing above (C) depicts the orientation of the GQ and triplex motifs (XR) in relation to the BPs. *** p < 0.0001 by one sample t-test. A) The number of deletion BPs associated with triplex motifs compared with reshuffled controls. Analysis including (left side) or excluding triplex-GQ hybrid motifs (right side). B) Same as (A) but with relaxed criteria for the detection of triplex motifs (min score=12) and GQ motifs (min score=26). C) The number of deletion BPs associated with GQ motifs compared with reshuffled controls. Relaxed settings (left side, min score=26) and default settings (right side, min score=47).

The six triplex motifs detected by the triplex package were significantly enriched around deletion breakpoints and when we excluded triplex-G-quadruplex hybrid motifs the result was attenuated but remained significant (**Fig. 3A**). Given the higher risk of spurious findings with only six motifs, we repeated the analysis using a relaxed definition of triplex and the results were fundamentally unchanged (**Fig. 3B**). Furthermore, our results were not sensitive to reasonable changes in the size of the search window around breakpoints (**Fig. S5A, B**), motif quality scores (**Fig. S5C, D**) or inclusion of overlapping motifs (**Fig. S5E-G**).

Analogous to the situation with MR motifs we tested if overlapping triplex-DR hybrid motifs could bias our results. Given the rarity of triplex motifs and the many DRs in the mitochondrial genome we choose an alternative approach rather than excluding triplex motifs that overlapped any DR half-site. We compared the fraction of triplex and G-quadruplex positive deletions associated with DRs (GQ^+^, DR^+^ and Trip^+^, DR^+^) and not associated with DRs (GQ^+^, DR^-^ and Trip^+^, DR^-^). We considered a deletion to be DR^+^ if both breakpoints were flanked by the same DR sequence. In this case, only 44% of Trip^+^ deletions associated with DRs whereas 66% of GQ^+^ deletions did (**Table S4**).

### Triplex forming motifs may be associated with mitochondrial disease breakpoints

Next, we sought to validate our findings on two recently published next generation sequencing datasets (**Hjelm et al. 2019, Persson et al. 2019**; **mtDNA Breakpoints**.**xlsx; Table S5**). We were able to confirm the enrichment of DR (**Fig. S6A, S7A**), MR (**Fig. S6A, S7A**) and G-quadruplex motifs (**Fig. 4A, B; S6C, D**) around deletion breakpoints. Additionally, we confirmed that hybrid MR-DR motifs are responsible in large part for the enrichment of MR motifs around breakpoints (**Fig. S6B, S7B**).

**Figure 4.**
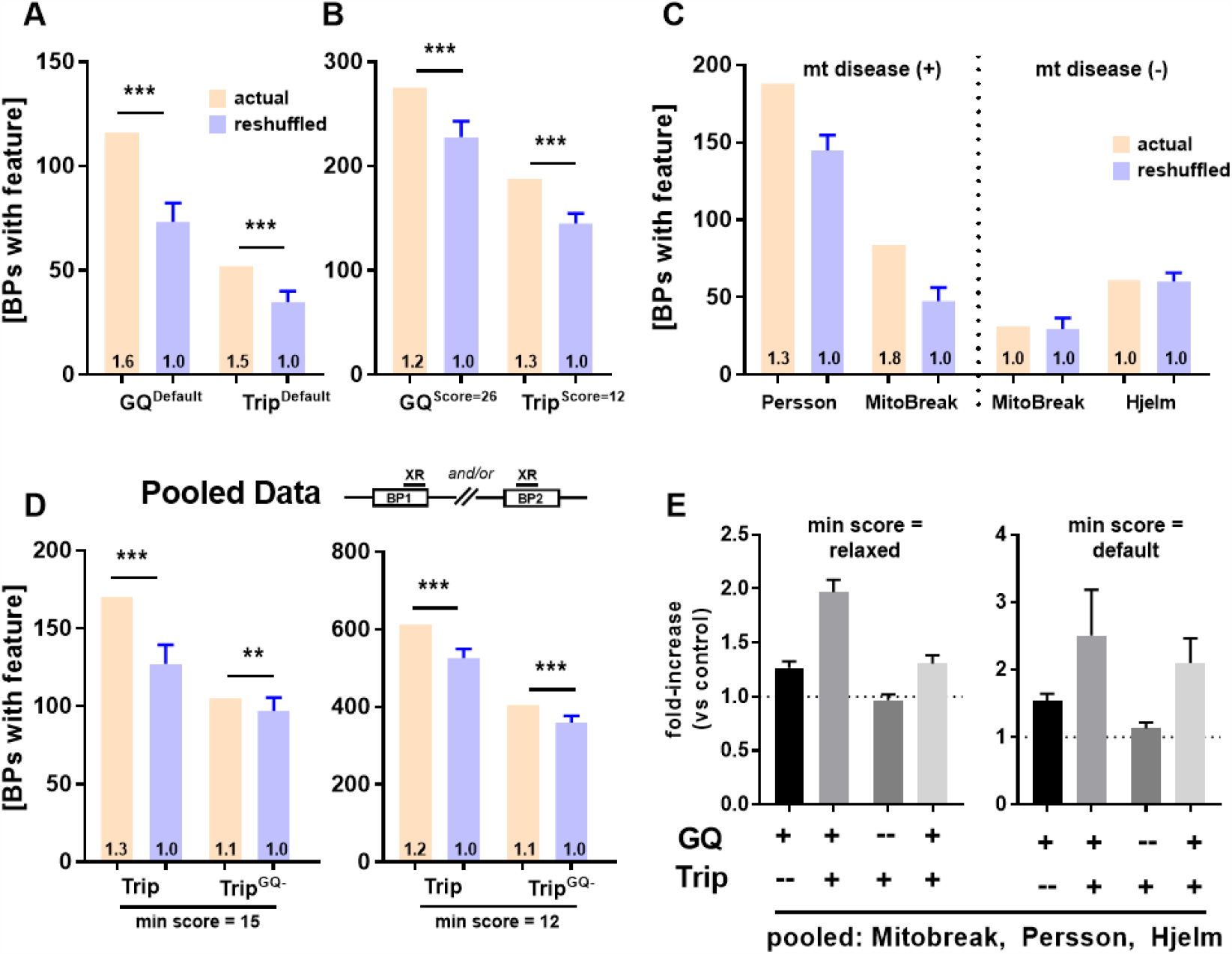
In the **Persson et al. (2019)** dataset, triplex and G-quadruplex (GQ) motifs are enriched around deletion breakpoints (BPs), using either default (A) or relaxed scoring criteria (B). Although triplex motifs predominate in mitochondrial disease datasets (C), we also find that triplex motifs are significantly enriched around BPs (D) after pooling the data from **MitoBreak, Persson et al. (2019) and Hjelm et al (2019)**. Finally, GQ and triplex motifs show stronger enrichment around BPs than either of them in isolation (E). Controls were generated by reshuffling the deletion BPs while maintaining their distribution (n=20, mean ±SD shown). The schematic drawing above (D) depicts the orientation of the motifs (XR) in relation to the BPs. *** p<0.0001, **p<0.001 by one sample t-test. A) The number of deletion BPs associated with GQ and triplex motifs compared with reshuffled controls (min score = default). B) The number of deletion BPs associated with GQ and triplex motifs compared with reshuffled controls (min score = relaxed). C) The number of deletion BPs associated with triplex motifs (relaxed settings, min score=12) stratified by mitochondrial disease status. MitoBreak data includes single and multiple mitochondrial deletion syndromes. D) The number of deletion BPs associated with triplex motifs, or with triplex motifs excluding triplex-GQ hybrid motifs (Trip^GQ-^), compared with reshuffled controls. Default settings (left side, min score=15) and relaxed settings (right side, min score=12). E) The fold-enrichment of GQ and triplex motifs around deletion BPs is shown. Motifs were considered overlapping if their midpoints were within 50 bp.

In contrast, we found that triplex motifs were not consistently enriched around breakpoints in the dataset of Hjelm et al. (**Fig. S6C, D**), which is based on post-mortem brain samples from patients without overt mitochondrial disease, whereas we saw enrichment in the dataset by Persson et al. (**Fig. 4A, B**), which is based on muscle biopsies from patients with mitochondrial disease. This unexpected discrepancy prompted us to take a second look at the MitoBreak data. In this dataset triplex motifs were significantly more enriched at breakpoints in the mtDNA single deletion subgroup compared to the healthy tissues subgroup (**Fig. S8**). In addition, we found more broadly that mitochondrial disease status might explain the heterogenous results across datasets we have seen (**Fig. 4C**).

Further strengthening our findings, triplex motifs were enriched in the MitoBreak and Persson et al. dataset regardless of the breakpoint shuffling method chosen and of our statistical assumptions (**Fig. S9**). What is more, triplex motifs were also enriched at breakpoints when we pooled all three datasets (**Fig. 4D**), although to a lesser extent.

Finally, G-quadruplex motifs close to triplex motifs were more strongly enriched at deletion breakpoints than solitary G-quadruplex motifs (**Fig. 4E; Fig. S10**), suggesting that triplex formation may further contribute to DNA instability.

### Repeats and lifespan: no support for the theory of resistant biomolecules

For our analysis, we focus on 11 bp long repeat motifs as short repeats are less likely to allow stable base pairing and longer repeats are rare (**Fig. S11**) and because results considering repeat motifs of different lengths usually agree with each other (**Table S6**; **Yang et al. 2013**). To allow comparability with other studies (**Lakshmanan et al. 2015**) we analyzed non D-loop motifs, but results for major arc motifs are numerically similar (**Table S7**).

First, consistent with **Yang et al. (2013)** we found that IR motifs show a negative correlation with the MLS of mammals in the unadjusted model. In addition, we identified ER motifs, a class of symmetrically related repeats, that show an even stronger inverse relationship with longevity (**Fig. 5A; Table 1**). However, these inverse correlations vanished after taking into account body mass, base composition and phylogeny in a PGLS model (**Table 1**). Second, in agreement with **Lakshmanan et al. (2015)** we found that DR motifs do not correlate with the MLS of mammals. The same was true for the symmetrically related MR motifs. Just as with IR motifs, modest inverse correlations vanished in the fully adjusted model (**Table 1**). We also found the same null results in two other vertebrate classes, birds and ray-finned fishes (**Table S6**). To gain hints as to causality, we finally tested if longer repeats, allowing more stable base pairing, show stronger correlations with MLS, but to our surprise we noticed the opposite (**Fig. S12A-D**).

**Figure 5.**
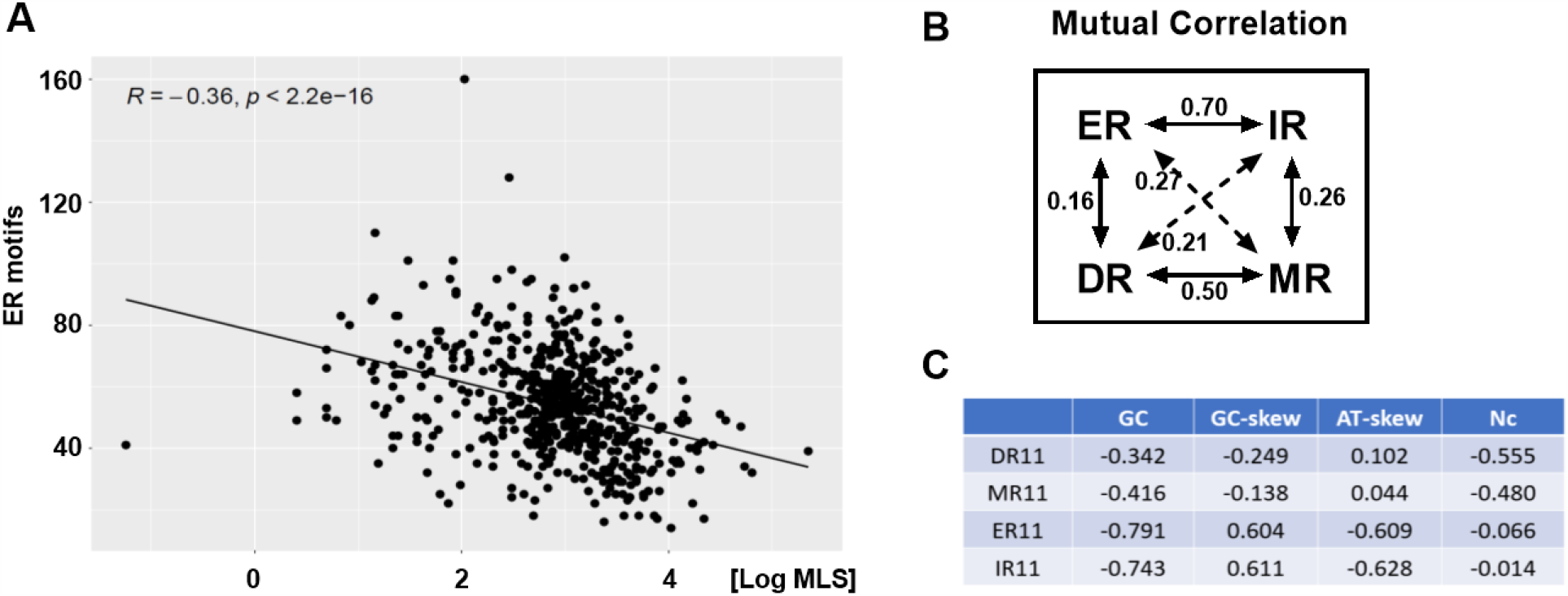
The number of everted repeat (ER) motifs is negatively correlated with species MLS in an unadjusted analysis (A). Repeats with a similar orientation correlate with each other (B). Direct repeat (DR) and mirror repeat (MR) motifs have a similar orientation since both half-sites are found on the same strand and in the case of ER and inverted repeat (IR) motifs the half-sites are on opposite strands. Finally, we show the major mtDNA compositional biases that co-vary with the four repeat classes (C) and may explain an apparent correlation with MLS. Data is for 11 bp long repeats and Pearson’s R is shown in (A-C).

**Table 1.** Correlation between potentially mutagenic motifs and species lifespan.

| Motif | Type | Raw | Adjusted |
| --- | --- | --- | --- |
| DR11 | 11bp | <u>-0.113</u> | 0.055 |
| MR11 | 11bp | <u>-0.155</u> | -0.002 |
| IR11 | 11bp | <u>-0.336</u> | <u>0.105</u> |
| ER11 | 11bp | <u>-0.356</u> | -0.047 |
| triplex | default | <u>-0.296</u> | <u>-0.211**</u> |
| triplex | relaxed | <u>-0.190</u> | <u>-0.127^</u> |
| GQ | default | <u>0.264</u> | 0.068 |
| GQ | relaxed | <u>0.283</u> | -0.097** |

The adjusted model takes into account body mass, GC content, GC skew, AT skew and number of effective codons. Significant correlations in the raw or adjusted model are bolded/underlined ( $p < 0.05$ ). The PGLS model additionally considers phylogeny. ^denotes p-values of $0.05 < p < 0.10$ in the PGLS model and \*\* p-values of $p < 0.05$ . The table shows Pearson's R.

Considering all four types of repeats together, we noticed that repeats with both half-sites on the same strand (DR and MR) or half-sites opposite strands (IR and ER) were correlated with each other (**Fig. 5B**) and with the same mtDNA compositional biases (**Fig. 5C**). Thus, for DR and MR motifs, an apparent relationship with MLS may be explained by their inverse relationship with GC content and for IR and ER motifs by an inverse relationship with GC content and a positive relationship with GC skew.

### Triplex motifs, not G-quadruplexes, show an inverse relationship with species lifespan

So far, no survey of G-quadruplex and triplex motifs has been conducted across species, although it is known that human mtDNA contains more G-quadruplexes than mouse, rat or monkey mtDNA (**Bharti et al. 2014**). First, we confirmed that different tools to detect G-quadruplex motifs agree with each other (**Lombardi and Londoño-Vallejo 2020; Fig. S13**). The results of different triplex detection tools, however, were inconsistent. While we were able to detect some overlap between the motifs found with Triplexator and the triplex package in human mtDNA (**Table S1**), we found that the two tools made very different predictions regarding triplex counts across species (**Fig. S14**).

The limited agreement between publicly available triplex detection tools raised the question whether our preferred tool, the triplex package in R, detects the same class of mutagenic triplex motifs as reported in earlier studies using in-house scripts (**Bacolla et al. 2016**). To answer this question, we reanalyzed a large dataset of over 500000 cancer associated breakpoints. In line with this earlier study, we found that both triplex and G-quadruplex motifs were significantly enriched around actual breakpoints compared to control breakpoints (**Fig. 6; Table S8**) and that breakpoints were preferably found in highly unstable regions with multiple such motifs (**Fig. S15**).

**Figure 6.**
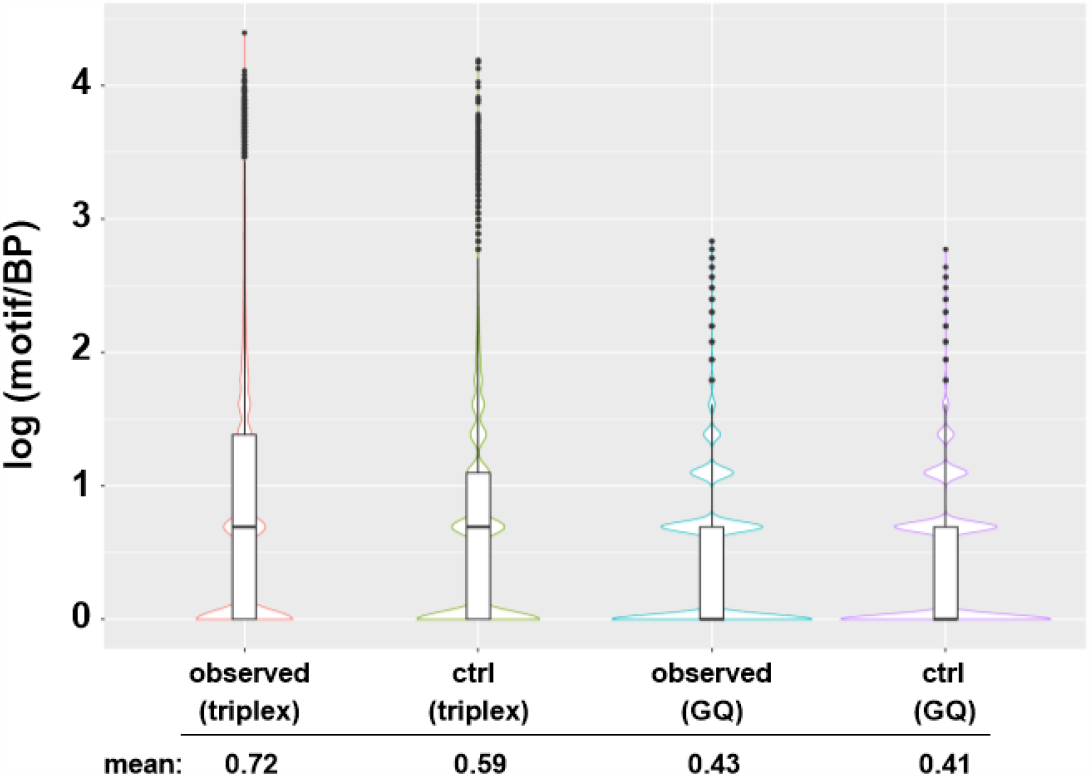
Actual cancer-related breakpoints (BPs) are associated with more triplex and G-quadruplex (GQ) motifs compared to control BPs (p < 2.2e-16; Wilcoxon rank sum test). To allow better visualization of the data, the number of motifs for each BP was log-transformed and log(0) values were excluded. Box whisker plots show the median, interquartile range and outliers while the underlying violin plot shows the actual distribution.

Next, we turned again to mtDNA to test whether mutagenic motifs are negatively correlated with MLS as predicted by the theory of resistant biomolecules. To the contrary, we found that G-quadruplex motifs were positively correlated with MLS (**Fig. S16**), although this may be secondary to their strong correlation with GC content. However, even after taking into account base composition and phylogeny using PGLS there was no convincing relationship between G-quadruplex motifs and MLS (**Table 1**).

In contrast, we found a moderate, negative correlation between intramolecular triplex motifs and the MLS of mammals (**Fig. 7A**) that was significant after correcting for body mass and base composition and, for triplex motifs using default scoring, also after correction for phylogeny using PGLS (**Table 1**). This result remained stable when we varied the score-cutoff and in fact triplexes were the only motif class for which we found that higher confidence motifs, i.e. motifs predicted to be more stable, showed a stronger relationship with MLS (**Fig. S12E**).

**Figure 7.**
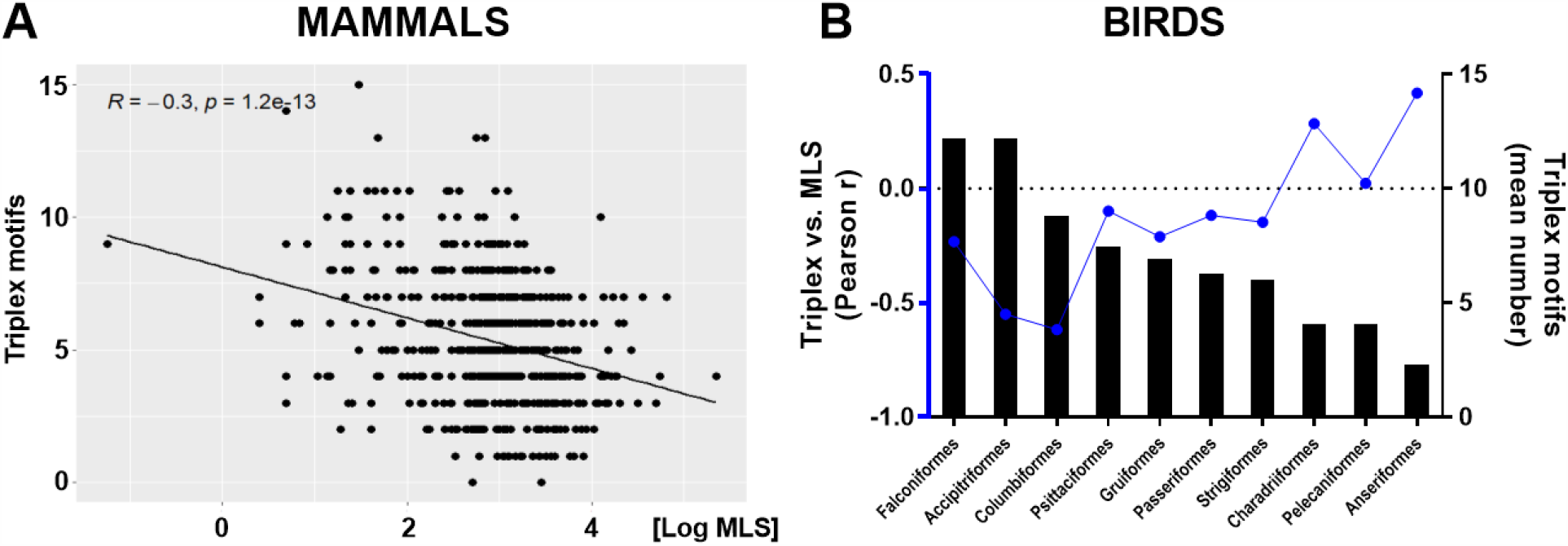
Across mammals, mitochondrial triplex motifs are inversely correlated with maximum lifespan (MLS) in an unadjusted analysis (A). Although, birds do not show a correlation between triplex motifs and MLS in the whole dataset (**Table S6**), the correlations between triplex motifs and MLS are stronger in bird orders with higher mean triplex levels (B). A) Mammals with a higher number of triplex motifs in their mtDNA (default settings) are on average shorter-lived. B) The higher the mean number of triplex motifs in a bird order (bar graphs) the stronger the inverse correlation between triplex motifs and MLS in the same order (blue line). All bird orders with more than 5 species in our dataset are included in this graph.

Although we found no significant relationship between triplex motifs and MLS of birds or ray-finned fishes, we noticed that this finding was modulated by the number of triplex motifs in the mtDNA of birds (**Fig. 7B**) and mammals (**Fig. S17B, D**). Phylogenetic orders with higher numbers of triplex motifs in their mtDNA showed a stronger inverse relationship between MLS and triplex counts than orders with few such motifs. Thus, when we split the bird and mammal data into orders with mitochondrial triplex levels above or below the median, we found a significant inverse relationship between MLS and triplex motifs in the high group for both birds and mammals (**Table S9**).

## DISCUSSION

### Repeat motifs and mtDNA deletions

Our goal was to define motifs associated with mtDNA deletions that would be then expected to show an inverse correlation with MLS under the theory of resistant biomolecules. We found that DR, MR, G-quadruplex and triplex motifs would be plausible candidates.

Specifically, our results support the consensus that DR and G-quadruplex motifs are major drivers of deletion mutagenesis (**Persson et al. 2019**) but not IR motifs (**Dong et al. 2014;** but see also: **Damas et al. 2012, Mikhailova et al. 2020**). In addition, we found a novel association between MR motifs and mtDNA deletions (**Fig. 2A**). This was unexpected since MR motifs do not allow for canonical base-pairing. Nonetheless, some studies have found MR motifs to be associated with indels (**Georgakopoulos-Soares et al. 2018**), perhaps due to their ability to form triplex DNA (**Kamat et al. 2016**).

Triplex forming motifs are mutagenic in bacteria (**Holder et al. 2015**), associated with deletions and translocations in cancer (**Bacolla et al. 2016**) and with mutations underlying various inherited diseases (**Kamat et al. 2016**), but little was known about their role in mitochondria. This is the first report that mitochondrial triplex motifs are indeed associated with mtDNA deletions (**Fig. 3**) and MLS (**Fig. 7**).

### Some mtDNA motifs colocalize and may interact

One key finding that emerged during our analysis was that motif classes correlate with each other, e.g., if both classes are enriched in DNA regions with a biased composition. Mitochondrial DR motifs are often close to MR and G-quadruplex motifs (**Fig. S18, Table S4**), IR motifs to ER motifs (**Table S10**) and triplex motifs to G-quadruplex motifs (**Fig. 3**). Our work is a first attempt to disentangle these interactions both in relation to deletion formation and to lifespan.

In principle one can correct for this by excluding hybrid motifs, i.e. two motifs located close to each other, from the analysis, as we have done in **Fig. 2-4**, but it is not always clear whether this is biologically sensible or overly conservative.

In the case of MR repeats we are inclined to conclude that they do not contribute to mtDNA instability, since there is little experimental data to implicate non-triplex-forming MRs in DNA instability and the enrichment of MR motifs at deletion breakpoints is attenuated after we exclude MR-DR hybrid motifs from the analysis (**Fig. 2C; Fig. S6, S7**).

In contrast, both triplex and G-quadruplex motifs are strongly implicated in DNA instability. A high G-content is known to promote G-quadruplex formation and to stabilize triplex DNA (**Buske et al. 2011, Kaufmann et al. 2019**) thus explaining their colocalization. Although G-quadruplexes may compete with the formation of triplex DNA, this is not always so (**Solé et al. 2017, del Mundo et al. 2017**).

Furthermore, in mtDNA, where overlapping triplex and G-quadruplex motifs can be located on opposite strands, a secondary structure on one strand could potentially promote formation of a secondary structure on the other strand by preventing reannealing of the two strands, as has been suggested for R-loops that promote the formation of G-quadruplexes (**De Magis et al. 2019**).

In support of the idea that motifs can interact, we found that G-quadruplex and triplex motifs in proximity to each other showed stronger associations with deletion breakpoints than isolated motifs, suggesting that their effects are additive (**Fig. S10**).

Similarly, it is not clear if the enrichment of G-quadruplex motifs at mtDNA deletion breakpoints needs to be corrected for their proximity to DR motifs (**Table S4**) found in regions of high GC-skew (**Fig. S18**). In a preliminary analysis we found that removal of DR-G-quadruplex hybrid sequences can greatly attenuate the over-representation of G-quadruplex motifs around breakpoints (data not shown). A thorough re-analysis of this phenomenon is beyond the scope of this work, but it does suggest that colocalization of different motifs is not a problem unique to our triplex dataset.

### mtDNA motifs in health and disease

We found that triplex motifs were particularly enriched around breakpoints in datasets from patients with mitochondrial disease (**Fig. 4C**). Conceivably, polymerase stalling associated with mitochondrial disease (**Wanrooij and Falkenberg 2010**) could allow more time for the formation of triplex DNA during replication, thereby explaining this finding. Consistent with this idea, a preprint by **Lakshmanan et al. (2018)** showed that mtDNA deletions in patients with mitochondrial disease due to POLG mutations are explained by predominantly short repeats and this bias towards short repeats was recently confirmed by a larger analysis that compared POLG patients to healthy controls (**Lujan et al. 2020**). These findings are intriguing because medium and long repeats are preferably associated with strand-slippage mechanisms whereas short repeats are consistent with strand breaks (**Nissanka et al. 2019, Lakshmanan et al. 2018**) which can be caused by ROS or secondary structures like G-quadruplex and triplex motifs.

### Analysis of mutagenic motifs and their relationship with lifespan casts doubt on the theory of resistant biomolecules

The theory of resistant biomolecules can be seen to imply a correlation between mutagenic motifs and MLS, but despite promising data from earlier studies (**Lakshmanan et al. 2012, Yang et al. 2013**), we found that neither G-quadruplex, DR, MR, IR nor ER motifs were associated with MLS after correcting for phylogeny and base composition (**Table 1**).

Perhaps the mtDNA is too streamlined to allow for the necessary changes in sequence. Given thousands of short, potentially mutagenic motifs (**Fig. S11**) that would need to be lost, we speculate it is more likely that nuclear proteins involved in mtDNA maintenance and metabolism account for the different rates of age-related deletion accumulation across species (**Guo et al. 2010**). In fact, repeats enable error prone DNA repair (**Tadi et al. 2016**), which might be advantageous under genotoxic stress to prevent wholescale mtDNA degradation and only detrimental late in life due to deletion accumulation, as would be predicted by the theory of antagonistic pleiotropy (**Campisi and Vijg 2009**).

The literature appears largely consistent with the null hypothesis. The work by **Yang et al. (2013)** may be considered the strongest case in favor of resistant biomolecules. They found an inverse relationship between IR motifs and MLS after correcting for phylogeny only. We found a similar correlation which, however, is fully accounted for by base composition. Two other findings of their paper are also consistent with the null hypothesis. Longer IR motifs, presumably allowing more stable base pairing, did not show a stronger inverse correlation with MLS, which we confirmed (**Fig. S12C**). Secondly, IR motifs with half-sites close together failed to show a stronger relationship with MLS, although these should form secondary structures more easily.

### The importance of mtDNA base composition in motif-lifespan correlations

It is striking that most of the findings were attenuated when we controlled for the base composition of mtDNA. Although the importance of such an adjustment was already highlighted by **Lakshmanan et al. (2015)** for DR motifs, we are the first to apply these corrections in the study of multiple DNA motifs.

We find it particularly informative to consider related motif classes together (**Fig. 5B, C**). ER motifs, for example, just like IR motifs show a strong correlation with GC content and GC skew, which in turn correlate with MLS. Importantly, ER motifs were not associated with mtDNA deletion breakpoints (**Fig. 2A, D**), are not known to be mutagenic, do not permit canonical base pairing and yet show a stronger inverse correlation with MLS than do IR motifs, which is fully attenuated after adjustment for base composition (**Table 1**). Thus, we conclude that most earlier results were driven by compositional bias.

### Can triplex motifs save the theory of resistant biomolecules?

We showed that intramolecular triplex motifs detected by the triplex package are inversely correlated with mammalian MLS after correcting for body mass, base composition and phylogeny (**Table 1**).

Although this moderate inverse correlation with MLS provides some support for this theory, it is not clear why major arc triplex motifs show a weaker than expected correlation with MLS (**Table S7**) nor if the triplex data can be generalized to non-mammalian species.

While we do not know a priori which motifs might be mutagenic in species with different mtDNA biology, we did find preliminary evidence that triplex motifs are inversely correlated with MLS in bird orders with high numbers of triplex motifs in their mtDNA (**Fig. 7B**) but no such correlation in ray-finned fishes. Since we observed the same trend in mammals, this suggests a potential threshold effect.

Perhaps a small number of triplex motifs in mtDNA is well tolerated, whereas larger numbers are progressively more destabilizing, especially if they occur in clusters as demonstrated for nuclear triplex motifs (**Fig. S15**).

### Going forward, finding the right motifs?

Why did an earlier study (**Oliveira et al. 2013**) fail to uncover an association between triplex motifs and mtDNA deletions? According to **Hon et al. (2013)** the nBMST tool used in that study only detects triplex-like mirror repeats and we also found it to be not very sensitive for the detection of putative triplex sequences. Given these shortcomings we also tested two other tools. The first one was Triplexator, which detects triplex target and triplex forming sites that could form both intra- and intermolecular triplexes, and the second one was the triplex package that detects intramolecular triplexes. We reasoned that intramolecular triplexes would be the most stable thus settling to use the latter tool. Using the triplex package we were able to confirm the well-established finding that nuclear triplex motifs are found near cancer-related breakpoints, supporting the validity of this tool (**Table S8**).

Although some of the sequences detected with Triplexator and the triplex package overlapped, many did not and in a comparison of 600 mammalian species the numbers of detected motifs did not correlate between the two tools (**Fig. S14**). This highlights a broader issue about the selection of motif detection tools. While multiple mature G-quadruplex detection tools exist that tend to agree with each other (**Fig. S13; Lombardi and Londoño-Vallejo 2020**), the choices for triplex motifs are more limited and there is little in vivo validation in mammalian genomes (**Kaufmann et al. 2019**).

Given these uncertainties, perhaps going forward we can refine the search parameters to find motifs that show a stronger correlation with lifespan and mtDNA deletions. To this end, our analysis of mitochondrial and nuclear breakpoints raises the possibility that hybrid and clustered motifs are particularly harmful (**Fig. 4E; Fig. S15**), but we have not yet analyzed these across genomes. Another issue that we did not address in detail is the type and orientation of triplex motifs involved in mtDNA deletions. Most importantly, future studies should explore to what extent these predicted intramolecular triplexes form in mitochondria.

## Supporting information

Species Data

mtDNA breakpoints

## ACKNOWLEDGEMENTS

We thank Doug Turnbull, Amy Vincent, David Meyer and Teresa Valencak for helpful comments and/or support. Reginald Smith for providing the python code to calculate Wright’s Nc, Mitya Toren for sharing updated MitoAge data and John A. Tainer for sharing triplex-related data.

## ABBREVIATIONS

BPs: mtDNA deletion break points
DR: direct repeats
ER: everted repeats
GQ: guanine-quadruplexes
IR: inverted repeats
MLS: species maximum lifespan
MR: mirror repeats
nBMST: non-B DNA motif search tool
Nc: number of effective codons
PGLS: phylogenetic generalized least squares
SD: standard deviation
Trip: Triplex forming motif
XR: any repeat half-site or motif
mtDNA: mitochondrial DNA

