## Supplementary Data for "Triplex and other DNA motifs show motif-specific associations with mitochondrial DNA deletions and species lifespan"

### SUPPLEMENTARY DATA - FIGURES

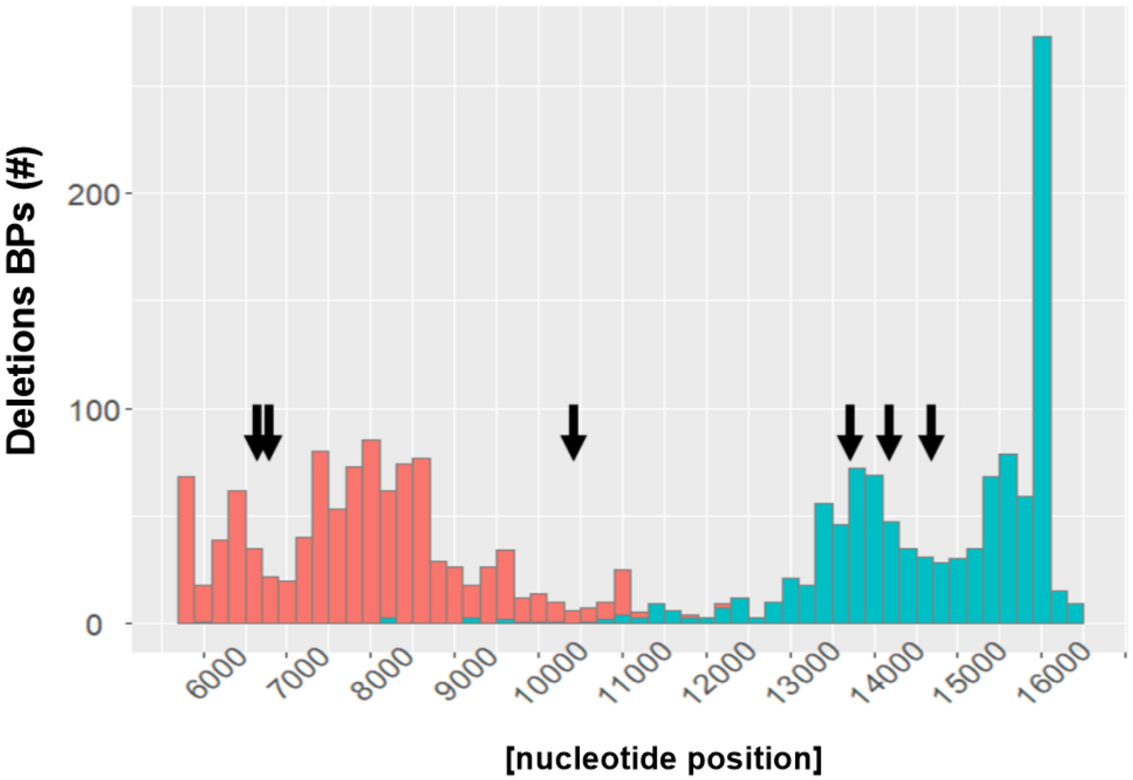

**Figure S1**
Distribution of breakpoints (BPs) inside the major arc with 5'-BPs shown in red, 3'-BPs shown in cyan and triplex motifs shown with arrows. Triplex motifs were detected using the triplex package in R with default scoring (min score=15). Breakpoints from the MitoBreak database.

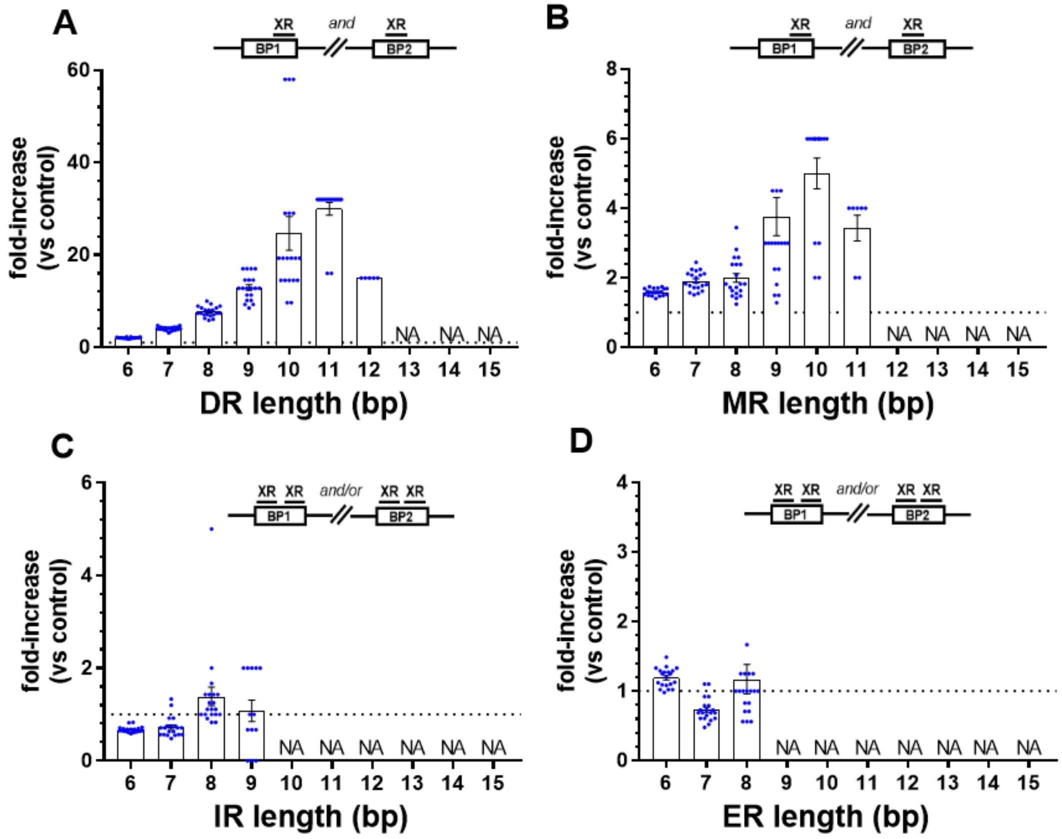

**Figure S2**

Direct repeat (DR; A) and mirror repeat (MR; B) motifs are enriched around actual breakpoints (BPs) compared to reshuffled breakpoints, but this is not true to the same extent for inverted repeats (IR; C) and everted repeats (ER; D). DR motifs show the strongest level of enrichment (note the y-axis). Controls were generated by reshuffling the deletion BPs while maintaining their distribution and the fold-change compared to controls was calculated (n=20, mean ±SD shown). The schematic drawings above (A-C) depict the orientation of the motifs (XR) in relation to the BPs. NA, analysis not possible due to limited sample size.

(A) DR motifs between 6 and 15 bps and their level of enrichment around deletion BPs.

(B) MR motifs between 6 and 15 bps and their level of enrichment around deletion BPs.

(C) IR motifs between 6 and 15 bps and their level of enrichment around deletion BPs.

(D) ER motifs between 6 and 15 bps and their level of enrichment around deletion BPs.

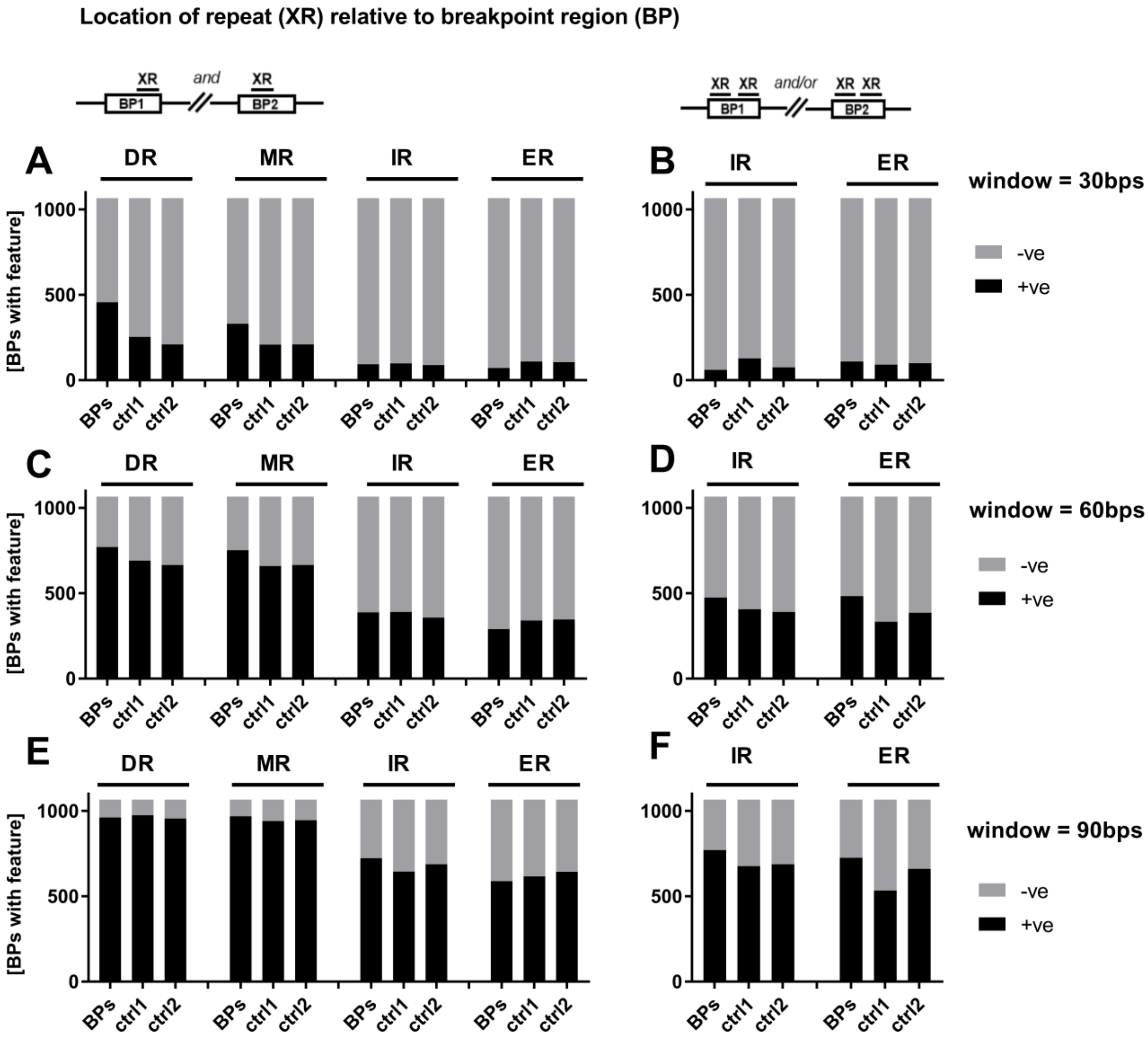

**Figure S3**

In this analysis we shifted each breakpoint (BP) by 200 bps to serve as its own control (ctrl1) and as a second control we generated fully random breakpoints within the major arc (ctrl2). Direct repeat (DR) and mirror repeat (MR) motifs remain somewhat enriched around actual BPs compared to reshuffled breakpoints even considering larger window sizes around BPs (A, C). In contrast, inverted repeat (IR) and everted repeat (ER) motifs are only enriched if we consider larger but not smaller window sizes (D, F) and the biological relevance of marginally increased IR motifs at BPs (+10 to 20%) is unclear.

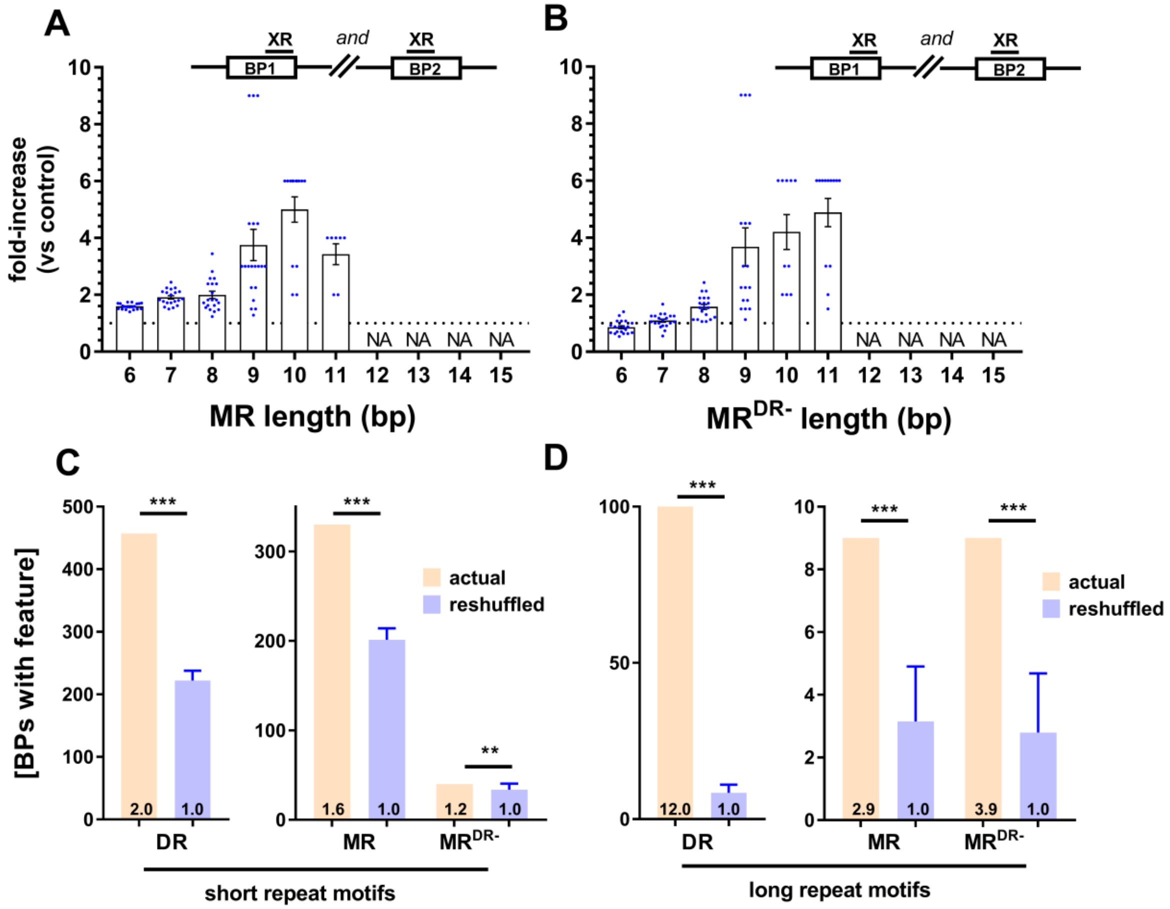

**Figure S4**

MR motifs of different lengths are significantly enriched around actual breakpoints (BPs) compared to reshuffled BPs (A, B). Removal of MR-DR hybrid motifs attenuates the correlation for short MR motifs but not for longer ones ($MR^{DR-}$; B). When we separate the motifs into two groups, 6 to 8 bp (C) and 9 to 15 bp long motifs (D), longer motifs are less abundant but more specifically enriched around BPs. Controls were generated by reshuffling the deletion BPs while maintaining their distribution (n=20, mean ±SD shown). The schematic drawings above (A, B) depict the orientation of the motifs (XR) in relation to the BPs. *** $p < 0.0001$, ** $p < 0.001$ by one sample t-test. Breakpoints from the MitoBreak database.

(A) MR motifs between 6 and 15 bps and their level of enrichment around deletion BPs.

(B) $MR^{DR-}$ motifs between 6 and 15 bps and their level of enrichment around deletion BPs.

(C) DR, MR and $MR^{DR-}$ motifs of 6 to 8 bp length and their level of enrichment around deletion BPs.

(D) DR, MR and $MR^{DR-}$ motifs of 9 to 15 bp length and their level of enrichment around deletion BPs.

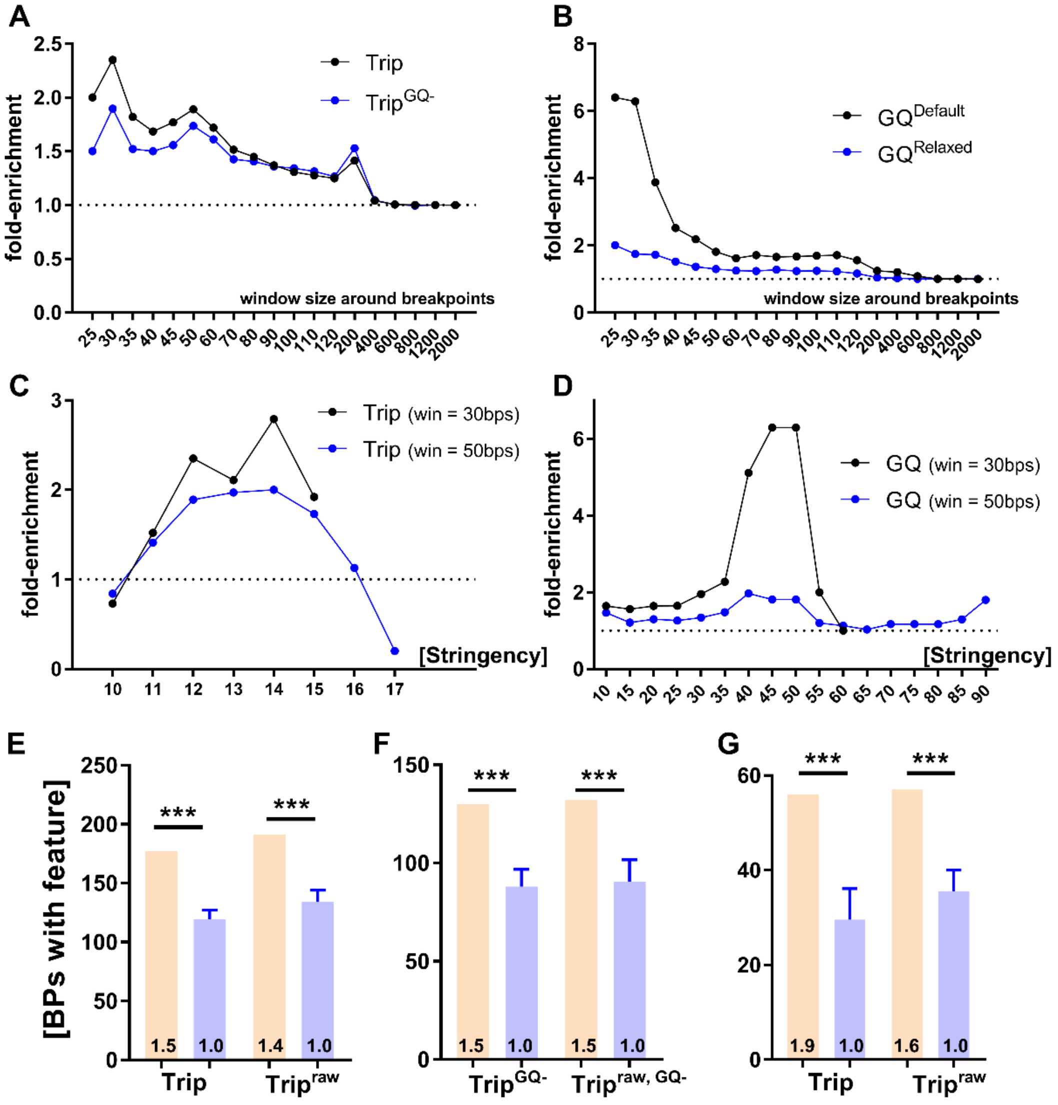

**Figure S5**

Triplex motifs (min score=12) are enriched around actual breakpoints (BPs) compared to shifted BPs across different window sizes (A) with similar results after removal of G-quadruplex (GQ)-triplex hybrid motifs ($Trip^{GQ-}$). GQ motifs are also enriched across different window sizes (B). In addition, it makes little difference if we vary the stringency (min score) of the detection algorithm. Both triplex (C) and GQ motifs (D) are enriched around actual BPs compared to shifted BPs across different min scores. Finally, in the main figure (Fig. 3) we only excluded predicted triplex motifs that were exact duplicates. Here, we show that "stringent" removal of both duplicate and partially overlapping motifs has only a marginal influence on the results (E-G), because these motifs usually overlap with the same BPs. *** $p < 0.0001$ by one sample t-test. Breakpoints from the MitoBreak database.

(A) Fold-enrichment of triplex motifs, and of triplex motifs excluding triplex-GQ hybrid motifs ($Trip^{GQ-}$), around actual BPs compared to shifted BPs (min score=12, relaxed settings). Different window sizes shown.

(B) Fold-enrichment of GQ motifs detected with default settings (min score = 47) or relaxed settings (min score = 26) around actual BPs compared to shifted BPs.
(C) Fold-enrichment of triplex motifs varying the minimum score cutoff, window sizes of 30 and 50 bps shown.
(D) Fold-enrichment of GQ motifs varying the minimum score cutoff, window sizes of 30 and 50 bps shown.
(E) Fold-enrichment after removal of all overlapping triplex motifs (n=22) compared with no removal (raw, n=33). Min score=12.
(F) Fold-enrichment after removal of all overlapping triplex motifs (n=15) compared with no removal (raw, n=33). Min score=12, excluding triplex-GQ hybrid motifs ($Trip^{GQ-}$)
(G) Fold-enrichment after removal of all overlapping triplex motifs compared with no removal (raw). Min score=15 (default).

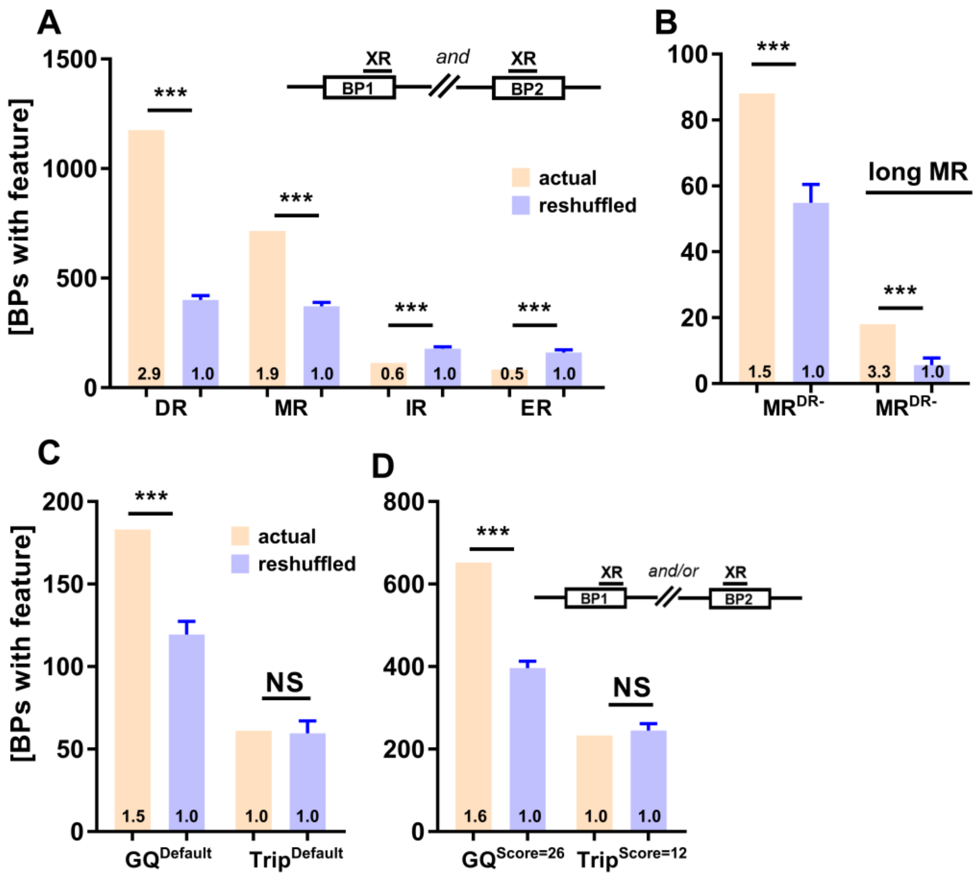

**Figure S6**
Our findings on the **Hjelm et al. (2019)** dataset agree with the MitoBreak data (**Fig. 2**) except for the relationship between triplex motifs and deletions (C, D). Direct repeat (DR) and mirror repeat (MR) motifs are significantly enriched around actual deletion breakpoints (BPs) compared to reshuffled BPs,

but the same is not true for inverted repeat (IR) and everted repeat (ER) motifs (A). The enrichment of MR motifs at deletion BPs is attenuated when MRs that have the same sequence as DR motifs are removed (B). GQ motifs are enriched at BPs, but the data for triplex motifs is inconsistent (C, D). Controls were generated by reshuffling the deletion BPs while maintaining their distribution (n=20, mean ±SD shown). *** p<0.0001 by one sample t-test.

(A) The number of deletion BPs associated with DR, MR, IR or ER motifs at both BPs compared with reshuffled controls.
(B) The number of BPs associated with MR motifs at both BPs after removal of hybrid MR-DR motifs ($MR^{DR-}$), compared with reshuffled controls. Long MR motifs have a length of 9 to 15 bps.
(C) The number of deletion BPs associated with GQ and triplex-forming motifs around BPs compared with reshuffled controls (min score = default).
(D) The number of deletion BPs associated with GQ and triplex-forming motifs around BPs compared with reshuffled controls (min score = relaxed).

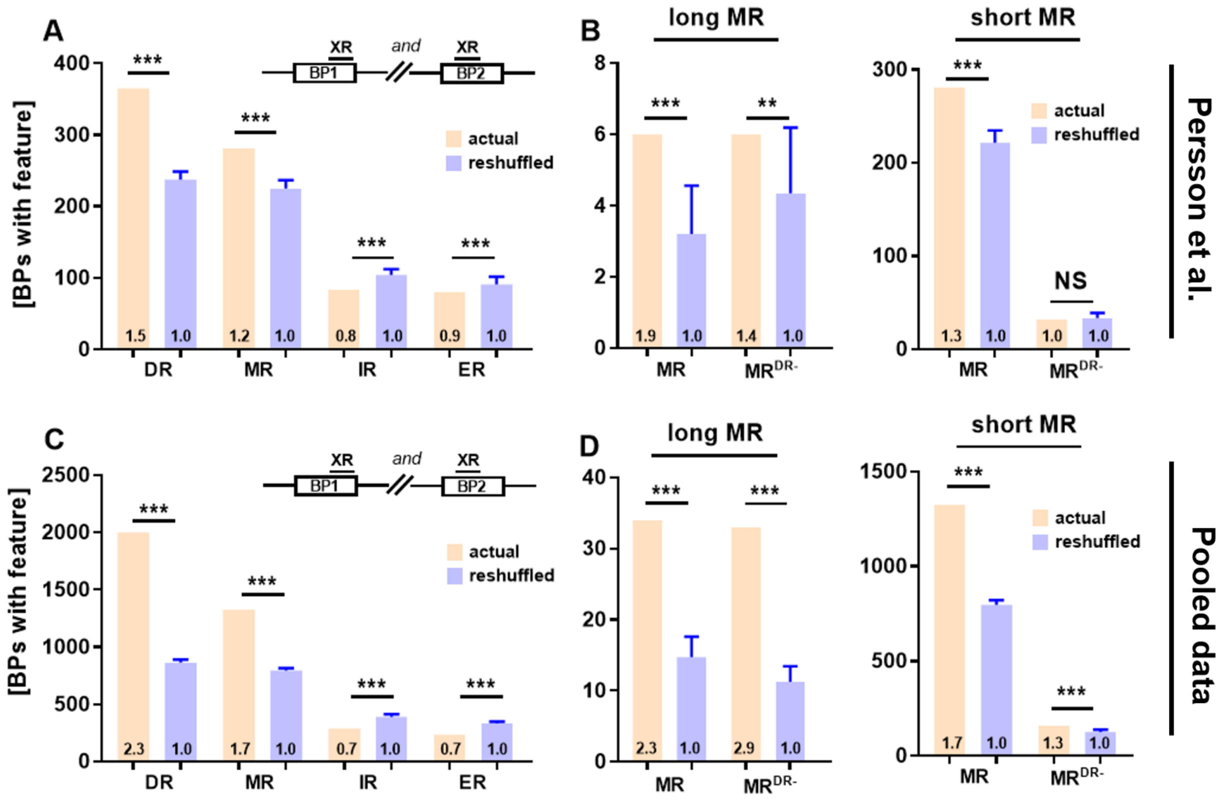

**Figure S7**
Our findings on the **Persson et al. (2019)** dataset and the pooled dataset (Hjelm + Persson + MitoBreak) agree with the MitoBreak data (**Fig. 2**). Direct repeat (DR) and mirror repeat (MR) motifs are significantly enriched around actual deletion breakpoints (BPs) compared to reshuffled BPs, but the same is not true for inverted repeat (IR) and everted repeat (ER) motifs (A, C). The correlation between MR motifs and deletion BPs is attenuated when adjusted for MR motifs that are equivalent to a DR motif (B, D). Controls were generated by reshuffling the deletion BPs while maintaining their distribution (n=20,

mean ±SD shown). The schematic drawing above (A, C) depicts the orientation of the repeat (XR) half-sites and motifs in relation to the BPs. *** p<0.0001, **p<0.001 by one sample t-test.

(A) The number of deletion BPs associated with DR, MR, IR or ER motifs at both BPs compared with reshuffled controls (based on **Persson et al. 2019**).
(B) The number of deletion BPs associated with MR motifs at both BPs, before (MR) and after removal of hybrid MR-DR motifs ($MR^{DR-}$), compared with reshuffled controls. Long MR motifs are defined as 9 to 15 bps and short MR motifs as 6 to 8 bps (based on **Persson et al. 2019**).
(C) Same as (A) but for the pooled dataset.
(D) Same as (B) but for the pooled dataset.

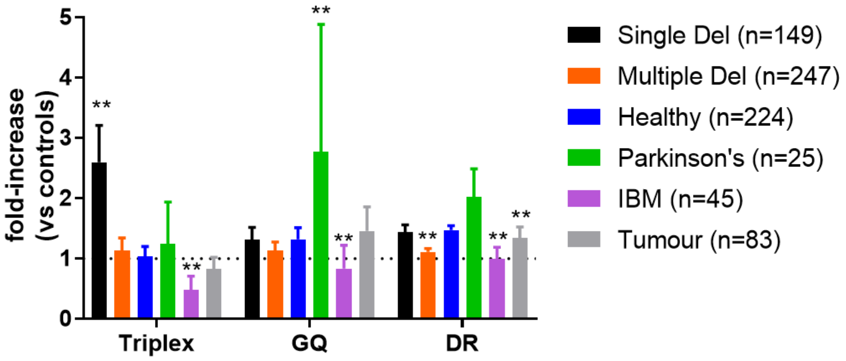

**Figure S8**
We split the MitoBreak data into subgroups according to disease etiology and for each subgroup we show fold-enrichment of motifs around actual breakpoints (BPs) compared to reshuffled BPs. The deletion BPs in the database comprise six groups: single mtDNA deletion syndromes, multiple mtDNA deletion syndromes, healthy tissues, Parkinson's disease, inclusion body myositis (IBM) or tumour tissues. We found that triplex motifs (min score = 12) are enriched compared to controls in several subgroups with the strongest increase in the single deletion group. This is different from the pattern seen for G-quadruplex (GQ; min score = 26) and direct repeat (DR) motifs.
Fold-enrichment calculated by comparison with n=20 reshuffled breakpoints (mean ±SD), window size around breakpoints = 50 bps for GQ and triplex and 30 bps for DR. Significance based on one-way ANOVA with Dunnett's post-hoc test, ** p < 0.05 compared with the healthy tissues group.

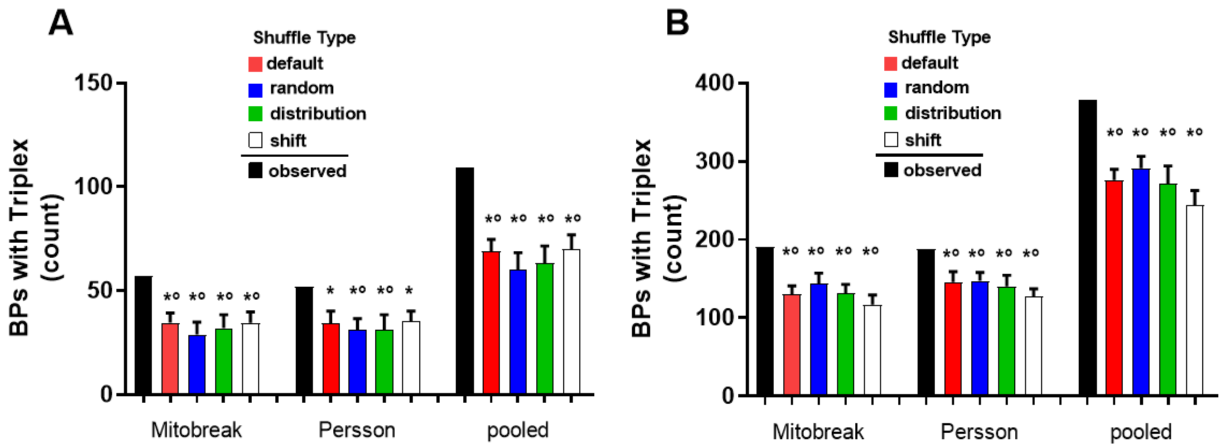

**Figure. S9**

The enrichment of triplex motifs around mtDNA deletion breakpoints (BPs) in the MitoBreak and Persson dataset is consistent when different BP shuffling methods are used and robust to statistical assumptions (one sample t-test vs. Fisher's exact test). "Default" shuffle, each mtDNA deletion as a whole is randomly redistributed within the major arc. "Random", individual mtDNA BPs are distributed randomly within the major arc. "Distribution", individual mtDNA BPs are redistributed in a way that approximates their original distribution. "Shift", each BP is randomly shifted by 100 to 300 bps, thereby maintaining the original distribution. Controls were generated by reshuffling the deletion BPs as described above (n=20, mean ±SD shown). * $p < 0.05$ by one sample t-test and ° $p < 0.05$ by Fisher's exact test.

(A) The number of observed BPs associated with triplex motifs compared to the number of reshuffled control BPs associated with triplex motifs (min score=15; default).
(B) Same as in (A) but with relaxed criteria for the detection of triplex motifs (min score=12).

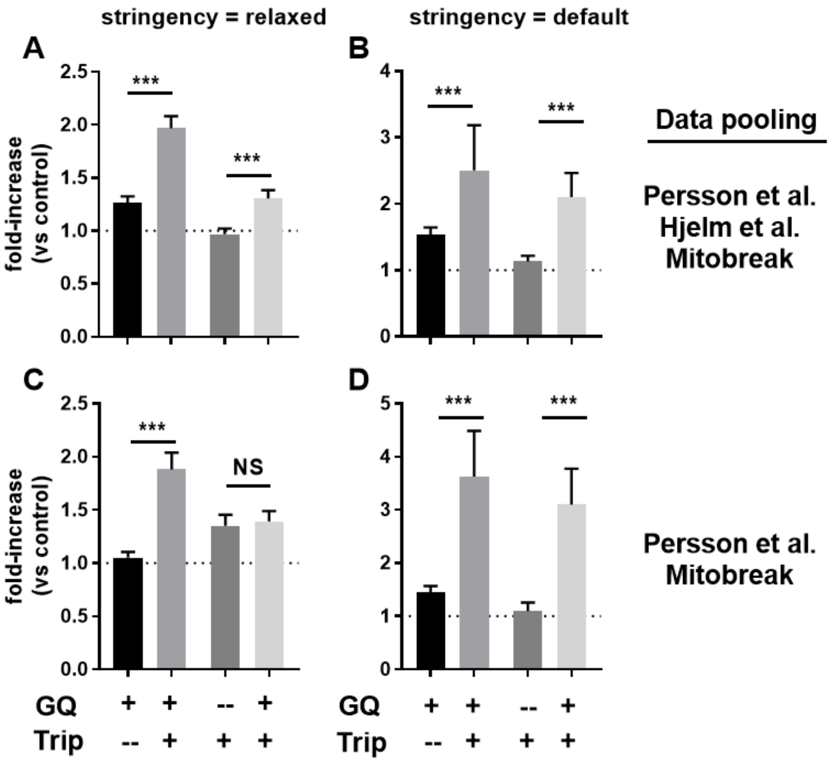

**Figure S10**

Fold-enrichment of motifs around actual breakpoints (BPs) compared to reshuffled breakpoints is shown. We define partially overlapping G-quadruplex (GQ)-triplex hybrid motifs if the sequence midpoints are within 50 bps of each other and we show that such hybrid motifs ($GQ^+$, $Trip^+$) are more strongly enriched around BPs compared to GQ ($GQ^+$, $Trip^-$) and triplex ($Trip^+$, $GQ^-$) motifs in isolation. Fold-enrichment calculated by comparison with n=20 reshuffled breakpoints (mean ±SD). *** $p < 0.0001$ by student's t-test.

(A) Fold-enrichment of the above motifs around mtDNA deletion BPs compared to reshuffled BPs. Data pooled from three studies, GQ min score = 26, Triplex min score = 12.
(B) Fold-enrichment of the above motifs around mtDNA deletion BPs compared to reshuffled BPs. Data pooled from three studies, GQ min score = 47, Triplex min score = 15.
(C) Fold-enrichment of the above motifs around mtDNA deletion BPs compared to reshuffled BPs. Data pooled from two studies, GQ min score = 26, Triplex min score = 15.
(D) Fold-enrichment of the above motifs around mtDNA deletion BPs compared to reshuffled BPs. Data pooled from two studies, GQ min score = 47, Triplex min score = 15.

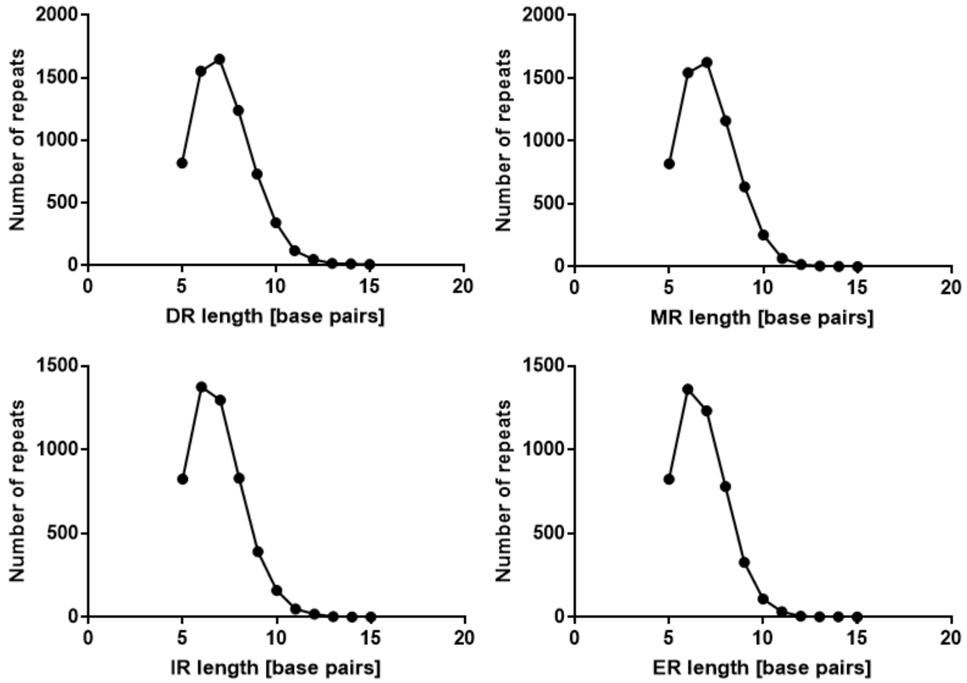

**Figure S11**
Long repeat motifs of any kind are rare in the human mitochondrial genome and their number decreases approximately in a log-linear fashion (not shown here). Since we plot unique repeats here, not counting multiple occurrences, the numbers for very short repeats decrease again due to their limited sequence diversity.

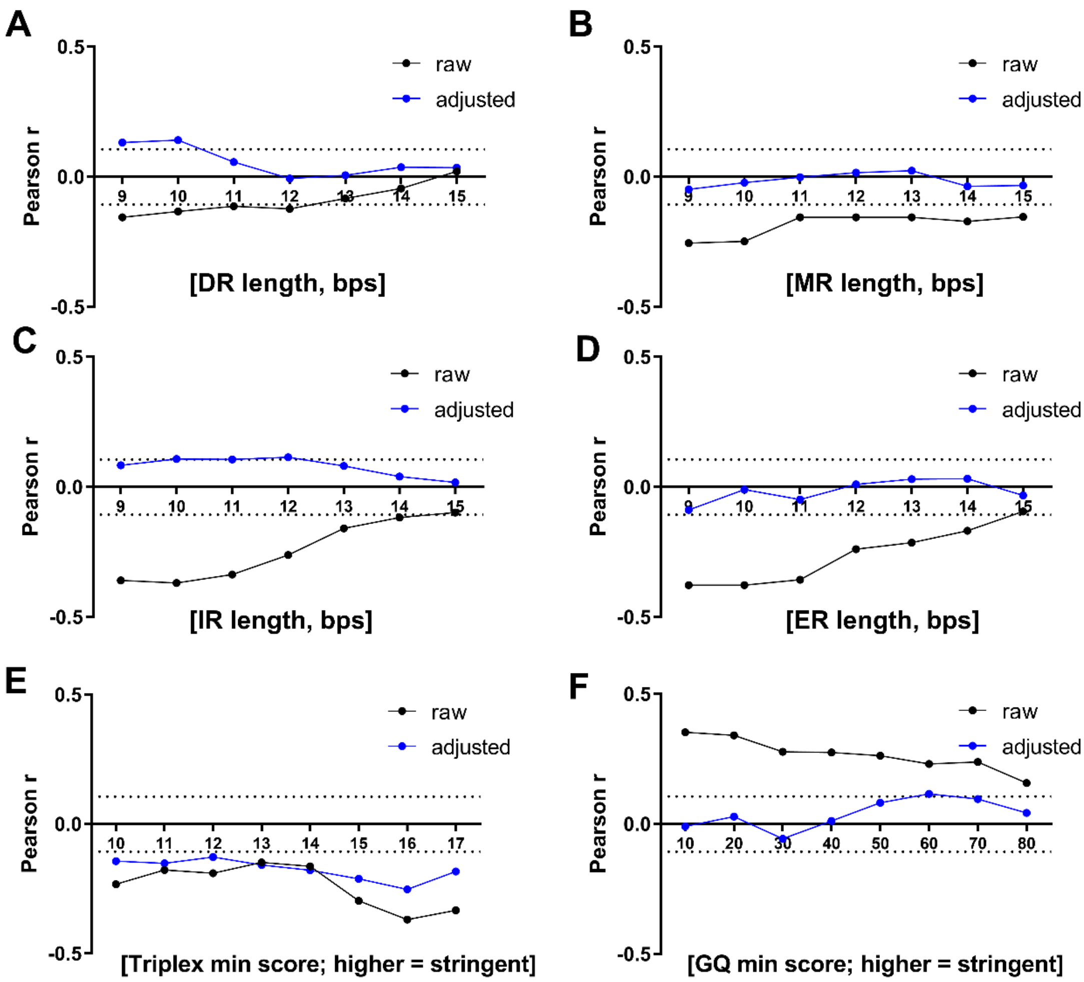

**Figure S12**

Secondary structures that are more thermodynamically stable would pose a larger threat to genome stability and should show a stronger inverse correlation with MLS. Although longer repeats will form more stable structures, they do not show a stronger correlation with MLS (A-D). Triplex (E) and G-quadruplex (GQ) motifs (F) with a higher minimum score should also be more stable, however, stable GQ motifs do not show an inverse correlation with MLS (F). Only for triplex motifs do we see the expected trend, a stronger inverse correlation with MLS in the case of more stable motifs (E). The critical R value at which p<0.01 is indicated.

(A) Longer direct repeat (DR) motifs do not consistently correlate with MLS after adjustment.

(B) Longer mirror repeat (MR) motifs do not consistently correlate with MLS after adjustment.

(C) Longer inverted repeat (IR) motifs do not consistently correlate with MLS after adjustment.

(E) Longer everted repeat (ER) motifs do not consistently correlate with MLS after adjustment.

(F) Triplex motifs show a consistent inverse correlation with MLS that is stronger for high scoring triplex motifs (i.e. motifs more likely to be stable).

(G) G-quadruplex motifs show a consistent positive correlation with MLS that is attenuated after adjustment. In contrast to triplex motifs, higher scoring GQ motifs (i.e. motifs more likely to be stable) do not show a stronger correlation with MLS.

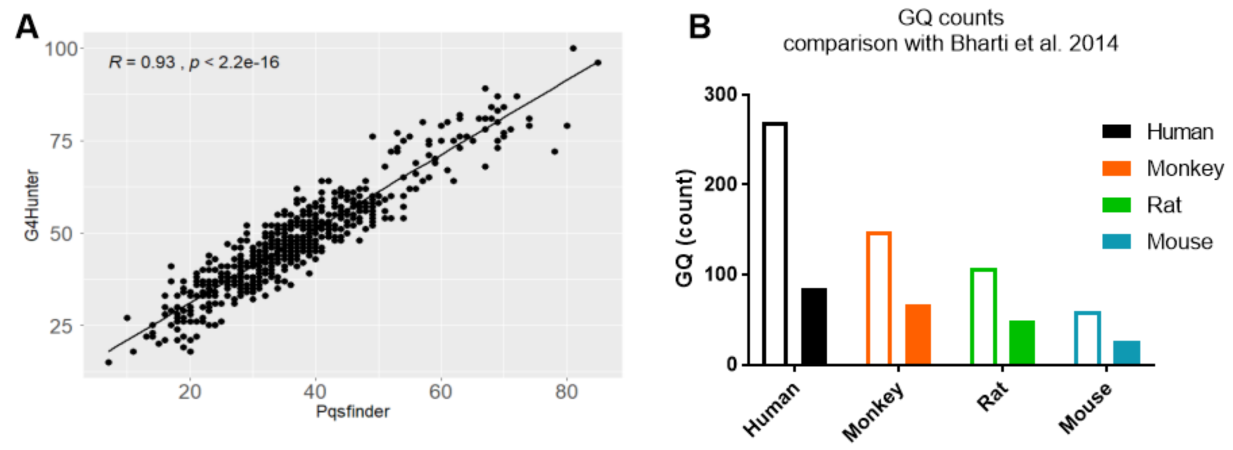

**Figure S13**
Two different G-quadruplex (GQ) prediction tools, pqsfinder and G4Hunter (both ran with default settings), perform similarly across 600 mammalian species (A). The data we generated using pqsfinder (filled bars) is comparable to Bharti et al. 2014 (open bars), although, our thresholds are more conservative yielding lower GQ counts (B). Our data is based on the mt genomes of NC_005943.1 (Macaca mulatta), AC_000022.2 (Rattus norvegicus), NC_005089.1 (Mus musculus) and NC_012920.1 (Homo sapiens).

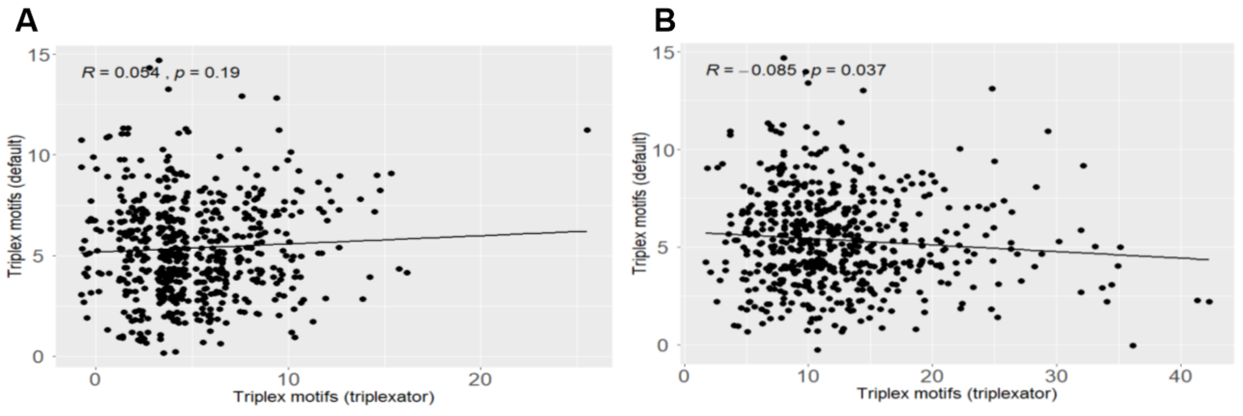

**Figure S14**
In contrast to the GQ prediction tools (Fig. S12), two different triplex motif prediction tools, the triplex package in R and triplexator (both ran with default settings), do not predict similar numbers of triplex motifs across 600 mammalian species. We use triplexator in -ds mode to detect potential triplex target sites (A) and in -ss mode to detect potential triplex forming oligonucleotides (B). Data shown as jitterplot for better visualization.

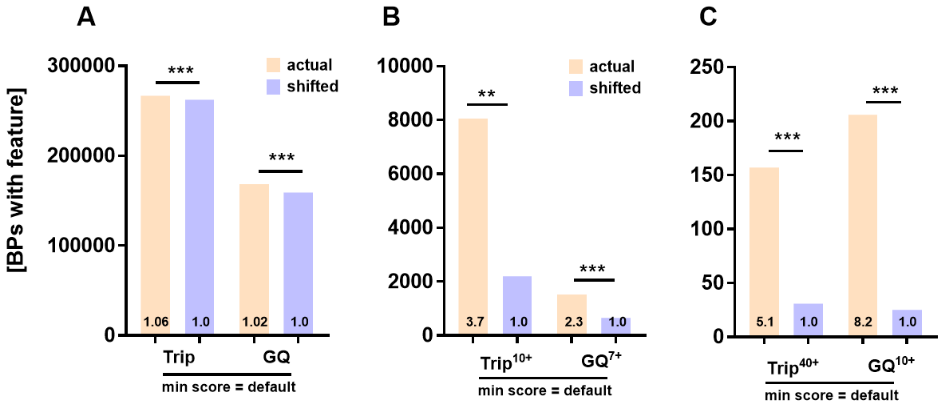

**Figure S15**

More actual, cancer-related breakpoints (BPs) compared to shifted control BPs are associated with triplex and G-quadruplex (GQ) motifs (A). The difference, however, is more pronounced when we consider highly unstable regions. More actual BPs compared to shifted control BPs are associated with multiple mutagenic motifs and thus lie in a highly unstable region (B, C; the number of mutagenic motifs is indicated by superscript). Each BP has a paired control and significance is determined by comparing the number of motifs per BP via Wilcoxon signed-rank test. ** p<0.01; *** p<0.001.

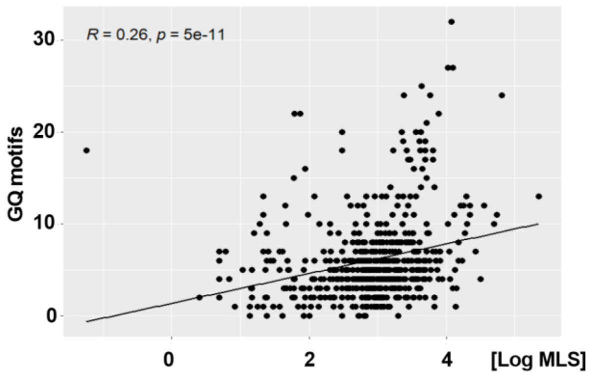

**Figure S16**

Across mammals, mitochondrial G-quadruplex (GQ) motifs show a positive correlation with species maximum lifespan (MLS) in an unadjusted analysis. GQ motifs were detected using default settings (min score = 47).

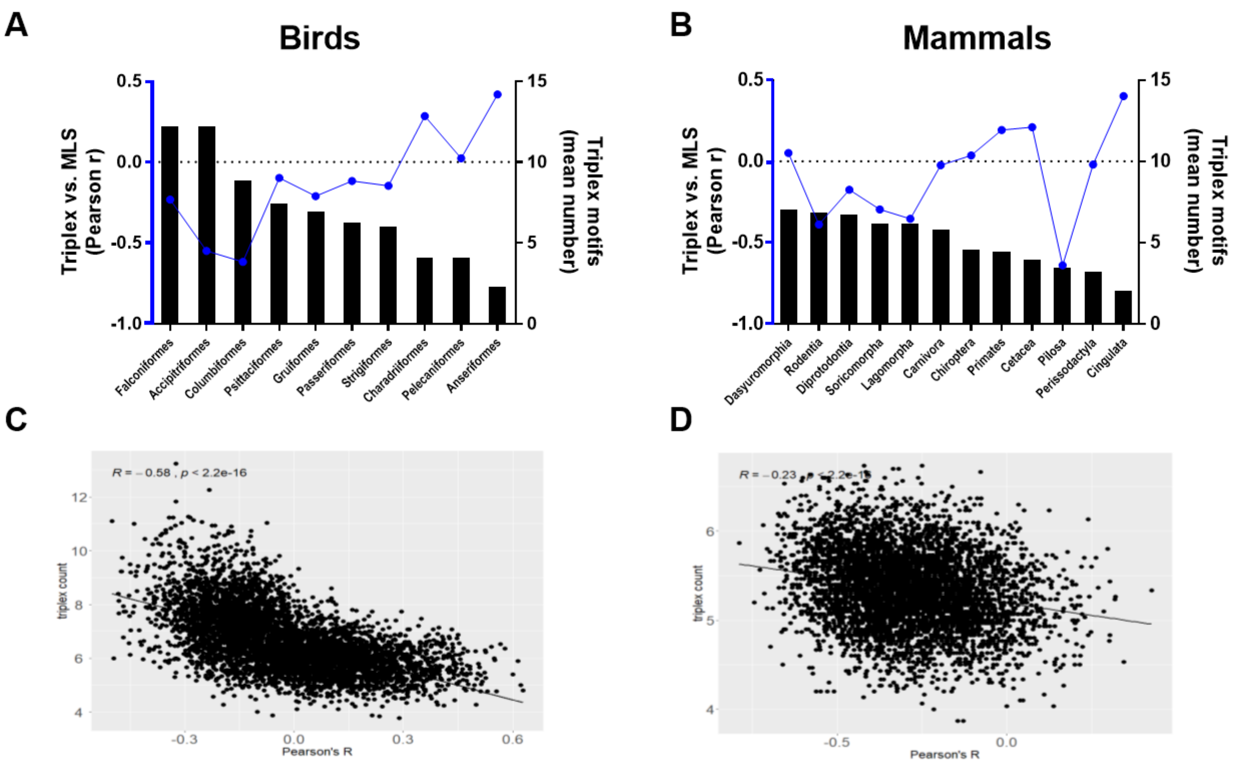

**Figure S17**

Phylogenetic orders (A, B) and randomly selected subgroups (C, D) of birds and mammals with a high number of triplex motifs in their mtDNA show a more robust inverse correlation between maximum lifespan (MLS) and triplex motifs, suggesting that levels of triplex motifs in mtDNA above a certain threshold could be detrimental to lifespan. Triplex motifs were detected using default settings (min score = 15).

A) The higher the mean number of triplex motifs in a bird order (bar graphs) the stronger the inverse correlation between triplex motifs and MLS in the same order (blue line). All bird orders with more than 5 species in our dataset are included in this graph.

(B) The higher the mean number of triplex motifs in a mammalian order (bar graphs) the stronger the inverse correlation between triplex motifs and MLS in the same order (blue line). All mammalian orders with more than 5 species in our dataset are included in this graph.

(C, D) We resampled 30 species from the bird (C) and mammal (D) dataset 5000 times. Then we calculated the correlation coefficient between triplex count and species MLS for each resampled subgroup. After this we plot the mean triplex count of each subgroup against the correlation coefficient of that subgroup.

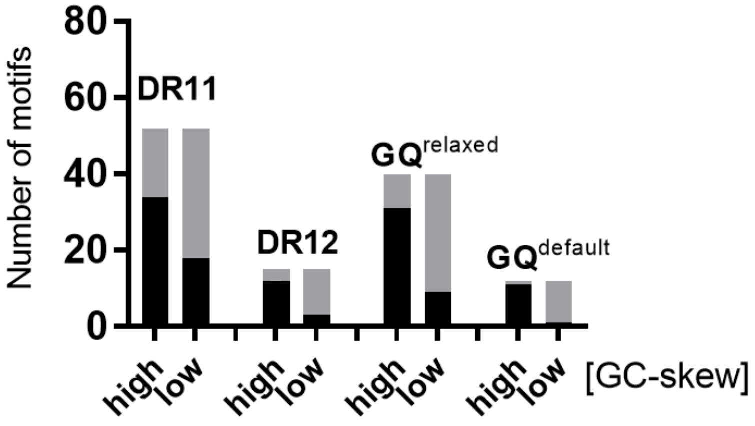

**Figure S18**

The human mtDNA was split into 100 bp windows and the number of direct repeat (DR) and G-quadruplex (GQ) motifs within highly and moderately GC-skewed windows is shown. Windows with a more negative skew than the median are defined as highly skewed and windows with a more positive skew are defined as low skew. We find that both DR and G-quadruplex motifs localize preferably in highly GC-skewed regions. DR11, DR12 … 11 and 12 bp long DR motifs. The dark shaded area shows the number of motifs associated with a given window and the grey shaded area the number of motifs not associated with a given window.

### SUPPLEMENTARY DATA - TABLES

Triplex package (default, min score = 15)

| | start | width | score | pvalue | ins | type | s | |
|---|---|---|---|---|---|---|---|---|
| [1] | 6676 | 26 | 17 | 7.3e-01 | 0 | 7 | + | [TAACTTACTACTCCGGAAAAAAAGAA] |
| [2] | 6689 | 31 | 15 | 9.9e-01 | 0 | 6 | + | [CGGAAAAAAAGAACCATTTGGATACATAGGT] |
| [3] | 10925 | 31 | 16 | 3.4e-01 | 0 | 1 | + | **[TTTTCCTCCGACCCCCTAACAACCCCCCTCC]** |
| [4] | 13751 | 20 | 15 | 6.2e-01 | 0 | 1 | + | **[TTTCCCCCGCATCCCCCTTC]** |
| [5] | 14097 | 19 | 15 | 6.2e-01 | 0 | 0 | + | **[CTTCCTCTCTTTCTTCTTC]** |
| [6] | 14599 | 21 | 15 | 6.2e-01 | 0 | 2 | - | **[TTTTCTTCTAAGCCTTCTCCT]** |

Triplex package (relaxed, min score = 12)

| | start | width | score | pvalue | ins | type | s | |
|---|---|---|---|---|---|---|---|---|
| [1] | 5830 | 25 | 13 | 1e+00 | 0 | 6 | + | [AAAAAGAGGCCTAACCCCTGTCTTT] |
| [2] | 6676 | 26 | 17 | 7.3e-01 | 0 | 7 | + | [TAACTTACTACTCCGGAAAAAAAGAA] |
| [3] | 6689 | 31 | 15 | 9.9e-01 | 0 | 6 | + | [CGGAAAAAAAGAACCATTTGGATACATAGGT] |
| [4] | 7434 | 19 | 14 | 1e+00 | 0 | 7 | + | **[ATAAAATCTAGACAAAAAA]** |
| [5] | 7465 | 19 | 12 | 1e+00 | 0 | 3 | - | [GAAACCAGCTTTGGGGGGT] |
| [6] | 7515 | 18 | 13 | 9.9e-01 | 0 | 2 | - | [TTTTTCTAATACCTTTTT] |
| [7] | 7515 | 18 | 12 | 1e+00 | 0 | 3 | - | [TTTTTCTAATACCTTTTT] |
| [8] | 8414 | 18 | 12 | 1e+00 | 0 | 0 | + | [CTCCTTACACTATTCCTC] |
| [9] | 8414 | 18 | 12 | 1e+00 | 0 | 1 | + | [CTCCTTACACTATTCCTC] |
| [10] | 9460 | 25 | 14 | 1e+00 | 0 | 5 | - | [AAAAAAACTTCTGAGGTAATAAATA] |
| [11] | 9477 | 27 | 14 | 8.9e-01 | 0 | 1 | + | [GTTTTTTTCTTCGCAGGATTTTTCTGA] |
| [12] | 9478 | 22 | 13 | 1e+00 | 0 | 4 | - | [AAAAATCCTGCGAAGAAAAAAA] |
| [13] | 10048 | 16 | 12 | 1e+00 | 0 | 6 | + | [AAAAAAGAGTAATAAA] |
| [14] | 10192 | 23 | 14 | 8.9e-01 | 0 | 0 | + | [CCCCCGCCCGCGTCCCTTTCTCC] |
| [15] | 10814 | 26 | 12 | 1e+00 | 0 | 6 | + | **[AAAAAACACATAATTTGAATCAACAC]** |

| | | | | | | | | |
|---|---|---|---|---|---|---|---|---|
| [16] | 10925 | 26 | 14 | 8.9e-01 | 0 | 2 | - | **[GGGGTTGTTAGGGGGTCGGAGGAAAA]** |
| [17] | 10925 | 31 | 16 | 3.4e-01 | 0 | 1 | + | **[TTTTCCTCCGACCCCCTAACAACCCCCCTCC]** |
| [18] | 10936 | 16 | 12 | 1e+00 | 0 | 4 | - | **[GGGGGTTGTTAGGGGG]** |
| [19] | 11032 | 25 | 13 | 1e+00 | 0 | 6 | + | [AAAAAAACTCTACCTCTCTATACTA] |
| [20] | 12106 | 29 | 12 | 1e+00 | 0 | 3 | - | **[AGGAAAACCCGGTAATGATGTCGGGGTTG]** |
| [21] | 12374 | 17 | 12 | 1e+00 | 0 | 1 | + | [CTTCCCTAATTCCCCCC] |
| [22] | 12383 | 26 | 14 | 8.9e-01 | 0 | 0 | + | [TTCCCCCCATCCTTACCACCCTCGTT] |
| [23] | 12396 | 30 | 13 | 1e+00 | 0 | 7 | + | **[TACCACCCTCGTTAACCCTAACAAAAAAAA]** |
| [24] | 12418 | 24 | 13 | 1e+00 | 0 | 6 | + | **[AAAAAAAACTCATACCCCCATTAT]** |
| [25] | 13218 | 20 | 13 | 1e+00 | 0 | 7 | + | [ACAAAATGACATCAAAAAAA] |
| [26] | 13751 | 20 | 15 | 6.2e-01 | 0 | 1 | + | **[TTTCCCCCGCATCCCCCTTC]** |
| [27] | 13752 | 18 | 14 | 8.9e-01 | 0 | 0 | + | **[TTCCCCCGCATCCCCCTT]** |
| [28] | 14097 | 19 | 15 | 6.2e-01 | 0 | 0 | + | **[CTTCCTCTCTTTCTTCTTC]** |
| [29] | 14492 | 19 | 12 | 1e+00 | 0 | 7 | + | [CCTAAATAAATTAAAAAAA] |
| [30] | 14504 | 27 | 13 | 1e+00 | 0 | 6 | + | [AAAAAAACTATTAAACCCATATAACCT] |
| [31] | 14599 | 21 | 15 | 6.2e-01 | 0 | 2 | - | **[TTTTCTTCTAAGCCTTCTCCT]** |
| [32] | 15443 | 18 | 13 | 9.9e-01 | 0 | 1 | + | **[CTTCTCTTCCTTCTCTCC]** |
| [33] | 16006 | 24 | 13 | 1e+00 | 0 | 5 | - | [AAAGAACAGAGAATAGTTTAAATT] |

Non-B DNA motif search tool (nBMST)

| type | start | end | width | sequence |
|---|---|---|---|---|
| Mirror_Repeat | 10077 | 10115 | 39 | ttaataatcaacaccctcctagccttactactaataatt |
| Mirror_Repeat | 11050 | 11111 | 62 | tatactaatctccctacaaatctccttaattataac(…) |

Triplexator (default, max error rate 5%)

| | Start | End | Score | Motif | Guanine.rate | TFO |
|---|---|---|---|---|---|---|
| 01 | 6213 | 6232 | 19 | Y | 0.55 | CTCTTACCTCCCTCTCTCCT |
| 02 | 6213 | 6232 | 19 | R | 0.55 | CTCTTACCTCCCTCTCTCCT |
| 03 | 6219 | 6238 | 19 | Y | 0.60 | CCTCCCTCTCTCCTACTCCT |
| 04 | 6219 | 6238 | 19 | R | 0.60 | CCTCCCTCTCTCCTACTCCT |
| 05 | 8803 | 8818 | 16 | M | 0.56 | ACACCAACCACCCAAC |
| 06 | 10831 | 10850 | 19 | M | 0.45 | **AATCAACACAACCACCCACA** |
| 07 | 10834 | 10853 | 19 | M | 0.55 | **CAACACAACCACCCACAGCC** |
| 08 | 12540 | 12560 | 20 | M | 0.48 | AGCCACAACCCAAACAACCCA |
| 09 | 12542 | 12562 | 20 | M | 0.52 | CCACAACCCAAACAACCCAGC |
| 10 | 14092 | 14117 | 25 | Y | 0.42 | **CTTTACTTCCTCTCTTTCTTCTTCCC** |
| 11 | 14092 | 14117 | 25 | R | 0.42 | **CTTTACTTCCTCTCTTTCTTCTTCCC** |
| 12 | 14097 | 14121 | 24 | Y | 0.48 | **CTTCCTCTCTTTCTTCTTCCCACTC** |
| 13 | 14097 | 14121 | 24 | R | 0.48 | **CTTCCTCTCTTTCTTCTTCCCACTC** |
| 14 | 14327 | 14347 | 20 | M | 0.57 | ACCACAACCACCACCCCATCA |
| 15 | 14613 | 14633 | 20 | M | 0.43 | **AAGAAAACCCCACAAACCCCA** |
| 16 | 15439 | 15462 | 23 | Y | 0.42 | CTTACTTCTCTTCCTTCTCTCCTT |
| 17 | 15439 | 15462 | 23 | R | 0.42 | CTTACTTCTCTTCCTTCTCTCCTT |
| 18 | 16277 | 16296 | 19 | M | 0.55 | ACCAACAAACCTACCCACCC |

Triplexator (relaxed, max error rate 7.5%)

| | Start | End | Score | Motif | Guanine.rate | TFO |
|---|---|---|---|---|---|---|
| 01 | 5931 | 5946 | 15 | M | 0.31 | ACAAACCACAAAGACA |
| 02 | 6213 | 6232 | 19 | Y | 0.55 | CTCTTACCTCCCTCTCTCCT |
| 03 | 6213 | 6232 | 19 | R | 0.55 | CTCTTACCTCCCTCTCTCCT |
| 04 | 6219 | 6238 | 19 | Y | 0.60 | CCTCCCTCTCTCCTACTCCT |
| 05 | 6219 | 6238 | 19 | R | 0.60 | CCTCCCTCTCTCCTACTCCT |
| 06 | 6486 | 6502 | 16 | Y | 0.53 | CTTCTCCTATCTCTCCC |
| 07 | 6486 | 6502 | 16 | R | 0.53 | CTTCTCCTATCTCTCCC |
| 08 | 6542 | 6557 | 15 | M | 0.56 | CAACCTCAACACCACC |
| 09 | 7397 | 7415 | 18 | M | 0.68 | CCCCCCACCCTACCACACA |
| 10 | 7443 | 7460 | 17 | R | 0.28 | **AGACAAAAAAGGAAGGAA** |
| 11 | 7443 | 7460 | 17 | Y | 0.28 | **AGACAAAAAAGGAAGGAA** |
| 12 | 7717 | 7733 | 16 | M | 0.35 | AACACTCACAACAAAAC |

| | | | | | | |
|---|---|---|---|---|---|---|
| 13 | 8272 | 8288 | 16 | Y | 0.71 | CCCCCTCTACCCCCTCT |
| 14 | 8272 | 8288 | 16 | R | 0.71 | CCCCCTCTACCCCCTCT |
| 15 | 8452 | 8468 | 16 | M | 0.41 | AAACACAAACTACCACC |
| 16 | 8597 | 8612 | 15 | Y | 0.44 | TTCTATTTCCCCCTCT |
| 17 | 8597 | 8612 | 15 | R | 0.44 | TTCTATTTCCCCCTCT |
| 18 | 8652 | 8667 | 15 | M | 0.44 | AATCACCACCCAACAA |
| 19 | 8803 | 8820 | 17 | M | 0.50 | ACACCAACCACCCAACTA |
| 20 | 9350 | 9366 | 16 | M | 0.41 | AACCAACACACTAACCA |
| 21 | 9664 | 9681 | 17 | M | 0.28 | AAAACAACCGAAACCAAA |
| 22 | 10267 | 10282 | 15 | Y | 0.56 | CCCTCCTTTTACCCCT |
| 23 | 10267 | 10282 | 15 | R | 0.56 | CCCTCCTTTTACCCCT |
| 24 | 10610 | 10625 | 15 | M | 0.56 | AACCCTCAACACCCAC |
| 25 | 10831 | 10850 | 19 | M | 0.45 | **AATCAACACAACCACCCACA** |
| 26 | 10834 | 10853 | 19 | M | 0.55 | **CAACACAACCACCCACAGCC** |
| 27 | 10886 | 10904 | 18 | M | 0.37 | AACCAAATCAACAACAACC |
| 28 | 10935 | 10952 | 17 | M | 0.67 | **ACCCCCTAACAACCCCCC** |
| 29 | 11227 | 11242 | 15 | Y | 0.62 | CTCCCTTCCCCTACTC |
| 30 | 11227 | 11242 | 15 | R | 0.62 | CTCCCTTCCCCTACTC |
| 31 | 12083 | 12099 | 16 | Y | 0.59 | TCCCCCATTCTCCTCCT |
| 32 | 12083 | 12099 | 16 | R | 0.59 | TCCCCCATTCTCCTCCT |
| 33 | 12090 | 12106 | 16 | Y | 0.53 | **TTCTCCTCCTATCCCTC** |
| 34 | 12090 | 12106 | 16 | R | 0.53 | **TTCTCCTCCTATCCCTC** |
| 35 | 12409 | 12426 | 17 | M | 0.28 | **AACCCTAACAAAAAAAAC** |
| 36 | 12540 | 12560 | 20 | M | 0.48 | AGCCACAACCCAAACAACCCA |
| 37 | 12542 | 12562 | 20 | M | 0.52 | CCACAACCCAAACAACCCAGC |
| 38 | 13290 | 13306 | 16 | M | 0.53 | CATCAACCAACCACACC |
| 39 | 13293 | 13308 | 15 | M | 0.50 | CAACCAACCACACCTA |
| 40 | 13770 | 13786 | 16 | M | 0.53 | **CCAAACAACAATCCCCC** |
| 41 | 14064 | 14079 | 15 | M | 0.44 | CACCTCAACCCAAAAA |
| 42 | 14092 | 14121 | 28 | Y | 0.43 | **CTTTACTTCCTCTCTTTCTTCTTCCCACTC** |
| 43 | 14092 | 14121 | 28 | R | 0.43 | **CTTTACTTCCTCTCTTTCTTCTTCCCACTC** |
| 44 | 14097 | 14126 | 28 | Y | 0.47 | **CTTCCTCTCTTTCTTCTTCCCACTCATCCT** |
| 45 | 14097 | 14126 | 28 | R | 0.47 | **CTTCCTCTCTTTCTTCTTCCCACTCATCCT** |
| 46 | 14181 | 14196 | 15 | M | 0.31 | ATACACCAACAAACAA |
| 47 | 14327 | 14347 | 20 | M | 0.57 | ACCACAACCACCACCCCATCA |
| 48 | 14387 | 14402 | 15 | M | 0.44 | AACCCCACTAAAACAC |
| 49 | 14551 | 14568 | 17 | M | 0.56 | AACACACCCGACCACACC |
| 50 | 14613 | 14633 | 20 | M | 0.43 | **AAGAAAACCCCACAAACCCCA** |
| 51 | 14639 | 14654 | 15 | M | 0.44 | AAACCCACACTCAACA |
| 52 | 14809 | 14825 | 16 | M | 0.65 | CCCCACCCCATCCAACA |
| 53 | 15439 | 15462 | 23 | Y | 0.42 | **CTTACTTCTCTTCCTTCTCTCCTT** |
| 54 | 15439 | 15462 | 23 | R | 0.42 | CTTACTTCTCTTCCTTCTCTCCTT |
| 55 | 15532 | 15548 | 16 | M | 0.59 | AAACACCCCTCCCCACA |
| 56 | 16162 | 16188 | 25 | M | 0.44 | AAAAACCCAATCCACATCAAAACCCCC |
| 57 | 16179 | 16194 | 15 | M | 0.62 | CAAAACCCCCTCCCCA |
| 58 | 16277 | 16296 | 19 | M | 0.55 | ACCAACAAACCTACCCACCC |

**Table S1. Triplex motifs detected by different motif prediction tools**

List of triplex motifs within the mtDNA major arc detected by the triplex package, the non-B DNA motif search tool (nBMST) and Triplexator. Before exclusion of redundant and overlapping motifs. Motifs found by Triplexator that overlap with motifs from the triplex package are underlined. The strand (s) is indicated for motifs found by the triplex package. Plus strand corresponds to the light-strand of mtDNA and minus strand to the heavy-strand.

| | | window size (bps) | | | | |
|---|---|---|---|---|---|---|
| **Motif** | **%error rate** | **20** | **30** | **40** | **50** | **60** |

| | | | | | | |
|---|---|---|---|---|---|---|
| triplexator | 5 | 1.50 | 1.86 | 1.31 | 1.33 | 1.24 |
| triplexator | 7.5 | 1.53 | 1.45 | 1.21 | 1.08 | 1.09 |
| triplexator | 10 | 1.10 | 1.33 | 1.27 | 1.13 | 1.10 |
| triplexator | 15 | 1.23 | 1.17 | 1.10 | 1.08 | 1.07 |
| rmGQ | 5 | 3.00 | 4.25 | 2.26 | 2.00 | 1.90 |

| **Motif** | **Stringency** | **20** | **30** | **40** | **50** | **60** |
|---|---|---|---|---|---|---|
| triplex | default | 6.00 | 1.92 | 1.62 | 1.73 | 1.65 |
| triplex | relaxed | 1.75 | 2.35 | 1.68 | 1.89 | 1.72 |
| rmGQ | default | 1.00 | 1.12 | 1.28 | 1.63 | 1.45 |
| rmGQ | relaxed | 2.00 | 3.14 | 2.05 | 2.21 | 1.84 |

| **Motif** | **Stringency** | **20** | **30** | **40** | **50** | **60** |
|---|---|---|---|---|---|---|
| GQ | default | 6.33 | 6.29 | 2.52 | 1.80 | 1.61 |
| GQ | relaxed | 2.52 | 1.80 | 1.51 | 1.29 | 1.25 |

**Table S2. Comparison of Triplexator, triplex package and pqsfinder data (fold-enrichment vs. control)**
This table shows the fold-enrichment of triplex motifs detected using two different methods (Triplexator and triplex package), and of G-quadruplex (GQ) motifs detected by the pqsfinder package, around actual deletion breakpoints (BPs) compared to a shifted control, where each BP was shifted by 200 bp towards the midpoint of the major arc. The acceptable maximal error rate (%) in Triplexator dictates the stringency of detection (higher = relaxed stringency). Analysis based on MitoBreak BPs.
rmGQ, overlapping GQ motifs removed.

```
Pqsfinder (default, min score = 47)

     start width score      sequence
 [1]  6290    30    52      [CCCTCCCTTAGCAGGGAACTACTCCCACCC]
 [2]  7397    16    53      [CCCCCCACCCTACCAC]
 [3]  7807    20    51      [CCTCATCGCCCTCCCATCCC]
 [4]  8262    34    67      [CCCTATAGCACCCCCTCTACCCCCTCTAGAGCCC]
 [5]  9243    28    55      [CCCAGCCCATGACCCCTAACAGGGGCCC]
 [6]  9526    49    85      [CCCCTACCCCCCAATTAGGAGGGCACTGGCCCCCAACAGGCATCA…]
 [7] 10184    24    60      [CCCTATATCCCCCGCCCGCGTCCC]
 [8] 10918    34    92      [CCCAACCTTTTCCTCCGACCCCCTAACAACCCCC]
 [9] 12084    32    67      [CCCCCATTCTCCTCCTATCCCTCAACCCCGAC]
[10] 12359    45    78      [CCACCCTAACCCTGACTTCCCTAATTCCCCCCATCCTTACCACCC]
[11] 13026    36    90      [CCCCTGACTCCCCTCAGCCATAGAAGGCCCCACCCC]
[12] 13647    49    62      [CCCCACCCTTACTAACATTAACGAAAATAACCCCACCCTACTAAAC…]
[13] 13755    31    61      [CCCCGCATCCCCCTTCCAAACAACAATCCCC]
[14] 14245    40    54      [CCCCGCACCAATAGGATCCTCCCGAATCAACCCTGACCCC]
[15] 14389    40    69      [CCCCACTAAAACACTCACCAAGACCTCAACCCCTGACCCC]
[16] 14771    47    49      [CCCCTAATAAAATTAATTAACCACTCATTCATCGACCTCCCCACCCC]
```

| | | | | |
|---|---|---|---|---|
| [17] | 15526 | 31 | 83 | [CCCCTTAAACACCCCTCCCCACATCAAGCCC] |
| [18] | 16159 | 35 | 51 | [CATAAAAACCCAATCCACATCAAAACCCCCTCCCC] |
| [19] | 16353 | 28 | 55 | [CCCTTCTCGTCCCCATGGATGACCCCCC] |

Pqsfinder (relaxed, min score = 26)

| | start | width | score | sequence |
|---|---|---|---|---|
| [1] | 6154 | 35 | 34 | [CCCTAATAATCGGTGCCCCCGATATGGCGTTTCCC] |
| [2] | 6257 | 15 | 27 | [GGAGGCCGGAGCAGG] |
| [3] | 6290 | 30 | 52 | [CCCTCCCTTAGCAGGGAACTACTCCCACCC] |
| [4] | 6421 | 48 | 43 | [CCCCTGCCATAACCCAATACCAAACGCCCCTCTTCGTCTGATCCG…] |
| [5] | 6539 | 49 | 40 | [CCGCAACCTCAACACCACCTTCTTCGACCCCGCCGGAGGAGGAGAC…] |
| [6] | 7095 | 46 | 29 | [CCCCTATTCTCAGGCTACACCCTAGACCAAACCTACGCCAAAATCC] |
| [7] | 7199 | 49 | 29 | [CCTATCCGGAATGCCCCGACGTTACTCGGACTACCCCGATGCATAC…] |
| [8] | 7397 | 16 | 53 | [CCCCCCACCCTACCAC] |
| [9] | 7466 | 28 | 38 | [CCCCCCAAAGCTGGTTTCAAGCCAACCC] |
| [10] | 7807 | 20 | 51 | [CCTCATCGCCCTCCCATCCC] |
| [11] | 8262 | 34 | 67 | [CCCTATAGCACCCCCTCTACCCCCTCTAGAGCCC] |
| [12] | 8369 | 41 | 42 | [CCCCAACTAAATACTACCGTATGGCCCACCATAATTACCCC] |
| [13] | 8464 | 12 | 32 | [CCACCTACCTCC] |
| [14] | 8559 | 50 | 41 | [CCCCACAATCCTAGGCCTACCCGCCGCAGTACTGATCATTCTATTT…] |
| [15] | 8619 | 10 | 35 | [CCCCACCTCC] |
| [16] | 8806 | 42 | 35 | [CCAACCACCCAACTATCTATAAACCTAGCCATGGCCATCCCC] |
| [17] | 8914 | 30 | 41 | [CCACAAGGCACACCTACACCCCTTATCCCC] |
| [18] | 9243 | 28 | 55 | [CCCAGCCCATGACCCCTAACAGGGGCCC] |
| [19] | 9289 | 14 | 29 | [CCTCCGGCCTAGCC] |
| [20] | 9413 | 15 | 27 | [CCACCACACACCACC] |
| [21] | 9526 | 49 | 85 | [CCCCTACCCCCCAATTAGGAGGGCACTGGCCCCCAACAGGCATCAC…] |
| [22] | 10184 | 24 | 60 | [CCCTATATCCCCCGCCCGCGTCCC] |
| [23] | 10267 | 27 | 42 | [CCCTCCTTTTACCCCTACCATGAGCCC] |
| [24] | 10612 | 26 | 42 | [CCCTCAACACCCACTCCCTCTTAGCC] |
| [25] | 10918 | 34 | 92 | [CCCAACCTTTTCCTCCGACCCCCTAACAACCCCC] |
| [26] | 10968 | 15 | 27 | [CCTGACTCCTACCCC] |
| [27] | 11130 | 33 | 26 | [CCACACTTATCCCCACCTTGGCTATCATCACCC] |
| [28] | 11206 | 31 | 39 | [CCTATTCTACACCCTAGTAGGCTCCCTTCCC] |
| [29] | 11407 | 25 | 43 | [CCCTAAAGCCCATGTCGAAGCCCCC] |
| [30] | 11532 | 12 | 32 | [CCTACCCCTTCC] |
| [31] | 11848 | 27 | 35 | [CCTCGCTAACCTCGCCTTACCCCCCAC] |
| [32] | 12084 | 32 | 67 | [CCCCCATTCTCCTCCTATCCCTCAACCCCGAC] |
| [33] | 12359 | 45 | 78 | [CCACCCTAACCCTGACTTCCCTAATTCCCCCCATCCTTACCACCC] |
| [34] | 12542 | 27 | 37 | [CCACAACCCAAACAACCCAGCTCTCCC] |
| [35] | 12927 | 45 | 33 | [CCCACAACAAATAGCCCTTCTAAACGCTAATCCAAGCCTCACCCC] |
| [36] | 13026 | 36 | 90 | [CCCCTGACTCCCCTCAGCCATAGAAGGCCCCACCCC] |
| [37] | 13119 | 13 | 30 | [CCGCTTCCACCCC] |
| [38] | 13647 | 49 | 62 | [CCCCACCCTTACTAACATTAACGAAAATAACCCCACCCTACTAAAC…] |
| [39] | 13755 | 31 | 61 | [CCCCGCATCCCCCTTCCAAACAACAATCCCC] |
| [40] | 13947 | 47 | 42 | [CCCCTATCTAGGCCTTCTTACGAGCCAAAACCTGCCCCTACTCCTCC] |
| [41] | 14043 | 16 | 26 | [CCAAATCTCCACCTCC] |
| [42] | 14115 | 23 | 28 | [CCCACTCATCCTAACCCTACTCC] |
| [43] | 14245 | 40 | 54 | [CCCCGCACCAATAGGATCCTCCCGAATCAACCCTGACCCC] |
| [44] | 14334 | 40 | 37 | [CCACCACCCCATCATACTCTTTCACCCACAGCACCAATCC] |
| [45] | 14389 | 40 | 69 | [CCCCACTAAAACACTCACCAAGACCTCAACCCCTGACCCC] |
| [46] | 14490 | 46 | 37 | [CCCCTAAATAAATTAAAAAAACTATTAAACCCATATAACCTCCCCC] |
| [47] | 14620 | 25 | 31 | [CCCCACAAACCCCATTACTAAACCC] |
| [48] | 14771 | 47 | 49 | [CCCCTAATAAAATTAATTAACCACTCATTCATCGACCTCCCCACCCC] |
| [49] | 14862 | 15 | 27 | [CCTGCCTGATCCTCC] |
| [50] | 15263 | 42 | 26 | [CCCACCCTCACACGATTCTTTACCTTTCACTTCATCTTGCCC] |
| [51] | 15368 | 48 | 28 | [CCCCTAGGAATCACCTCCCATTCCGATAAAATCACCTTCCACCCT…] |
| [52] | 15526 | 31 | 83 | [CCCCTTAAACACCCCTCCCCACATCAAGCCC] |
| [53] | 15658 | 11 | 33 | [CCCCATCCTCC] |
| [54] | 16159 | 35 | 51 | [CATAAAAACCCAATCCACATCAAAACCCCCTCCCC] |
| [55] | 16260 | 37 | 44 | [CCCTCACCCACTAGGATACCAACAAACCTACCCACCC] |
| [56] | 16353 | 28 | 55 | [CCCTTCTCGTCCCCATGGATGACCCCCC] |

```
[57] 16400    12    32      [CCACCATCCTCC]
```

**Table S3. G-quadruplex motifs in the major arc of mtDNA**
List of G-quadruplex (GQ) motifs detected by pqsfinder. All GQ motifs but one were found on the heavy-strand (motif #2).

| Motif | $DR^+$ | $DR^-$ | ratio ($DR^+$ vs $DR^-$) | p-value |
|---|---|---|---|---|
| $GQ^+$ (default) | 82 | 42 | 2.0 | $p<0.05$ |
| $Triplex^+$ (default) | 28 | 30 | 0.9 | |
| $GQ^+$ (relaxed) | 189 | 114 | 1.7 | $p<0.0001$ |
| $Triplex^+$ (relaxed) | 86 | 119 | 0.7 | |

**Table S4. Colocalization of G-quadruplex and triplex motifs with DR motifs**
G-quadruplex (GQ) motifs preferably colocalize with deletions that are associated with direct repeat motifs (DR), whereas triplex motifs do not. Significance based on Fisher's exact test. Deletion data from the MitoBreak database.

| | 5'-Breakpoints | | 3'-Breakpoints | | deletion | | |
|---|---|---|---|---|---|---|---|
| | median | mean | median | mean | median size (bp) | major arc deletions (N) | subjects (N) |
| MitoBreak | 7962 | 8077 | 15149 | 14741 | 7122 | 1066 | NA |
| Persson | 8396 | 8708 | 15063 | 14571 | 6125 | 1114 | 5 |
| Hjelm | 8106 | 8446 | 14341 | 14075 | 5926 | 1894 | 93 |

**Table S5. Comparison of mtDNA deletion breakpoint datasets**
Breakpoint positions and other characteristics of the three datasets we used (**MitoBreak, Persson et al. 2019 and Hjelm et al 2019).**

| | | Mammals | | Birds | | Ray-finned fishes | |
|---|---|---|---|---|---|---|---|
| Motif | Type | Raw | Adjusted | Raw | Adjusted | Raw | Adjusted |
| DR11 | 11bp | **-0.113** | 0.055 | 0.090 | **0.140** | 0.117 | -0.026 |
| MR11 | 11bp | **-0.155** | -0.002 | 0.037 | 0.115 | 0.065 | -0.003 |
| IR11 | 11bp | **-0.336** | **0.105** | -0.125 | -0.016 | 0.086 | **0.250** |
| ER11 | 11bp | **-0.356** | -0.047 | -0.073 | 0.002 | -0.020 | 0.087 |

| | | | | | | | |
|---|---|---|---|---|---|---|---|
| DR | 9 to 15bp | **-0.149** | **0.127** | 0.059 | 0.107 | **0.231** | **0.212** |
| MR | 9 to 15bp | **-0.255** | -0.025^ | 0.068 | **0.167** | 0.140 | 0.053 |
| IR | 9 to 15bp | **-0.369** | -0.058 | -0.012 | 0.140 | -0.032 | 0.113 |
| ER | 9 to 15bp | **-0.384** | **0.134** | -0.074 | 0.052 | -0.093 | **0.212** |
| triplex | default | **-0.296** | **-0.211**** | -0.073 | 0.003 | -0.083 | 0.040 |
| triplex | relaxed | **-0.190** | **-0.127^** | -0.012 | 0.077 | 0.091 | **0.354**** |
| GQ | default | **0.264** | 0.068 | 0.108 | **0.174** | 0.017 | **0.292^** |
| GQ | relaxed | **0.283** | -0.097** | 0.042 | 0.119 | -0.020 | **0.269** |

**Table S6. Correlation between motifs and MLS in birds and ray-finned fishes (actinopterygii)**
The adjusted model takes into account GC content, GC skew, AT skew and number of effective codons. Significant correlations in the raw or adjusted model are bolded/underlined (p<0.05). The PGLS model additionally considers phylogeny. ^denotes p-values of 0.05<p<0.10 in the PGLS model and ** p-values of p<0.05. The table shows Pearson's R.

| | | non-D loop mtDNA | | major arc mtDNA | |
|---|---|---|---|---|---|
| **Motif** | **Type** | **Raw** | **Adjusted** | **Raw** | **Adjusted** |
| DR11 | 11bp | **-0.113** | 0.055 | **-0.162** | 0.052 |
| MR11 | 11bp | **-0.155** | -0.002 | **-0.126** | -0.006 |
| IR11 | 11bp | **-0.336** | **0.105** | **-0.314** | 0.043 |
| ER11 | 11bp | **-0.356** | -0.047 | **-0.313** | -0.046 |
| triplex | default | **-0.296** | **-0.211**** | **-0.240** | **-0.127^** |
| triplex | relaxed | **-0.190** | **-0.127^** | **-0.135** | -0.059 |
| GQ | default | **0.264** | 0.068 | **0.263** | 0.057 |
| GQ | relaxed | **0.283** | -0.097** | **0.278** | **-0.107**** |

**Table S7. Correlation between potentially mutagenic motifs in the mtDNA and species lifespan**
The adjusted model takes into account body mass, GC content, GC skew, AT skew and number of effective codons. Significant correlations in the raw and adjusted model are bolded/underlined ($p<0.05$). The PGLS model additionally considers phylogeny. ^denotes p-values of $0.05<p<0.10$ in the PGLS model and ** p-values of $p<0.05$. The table shows Pearson's R.

| | **Triplex motifs** | | | **GQ motifs** | | |
|---|---|---|---|---|---|---|
| | **actual** | **control** | **fold-enrichment** | **actual** | **control** | **fold-enrichment** |
| **mean (motifs/BP)** | 0.948 | 0.674 | 1.41 | 0.822 | 0.785 | 1.05 |
| **pseudo-median** | 2 | 1.999942 | NA | 1.500065 | 1.500044 | NA |
| **CI max** | 2.000036 | 1.999936 | NA | 1.500029 | 1.500063 | NA |
| **CI min** | 2.000002 | 1.999948 | NA | 1.500006 | 1.50004 | NA |
| **Total BPs** | 577994 | 577994 | NA | 577994 | 577994 | NA |
| **BPs with X motifs:** | | | | | | |
| 0 | 408279 | 418334 | 0.98 | 311129 | 315643 | 0.99 |
| 1 | 77810 | 79537 | 0.98 | 143692 | 144915 | 0.99 |
| 2 | 33749 | 34258 | 0.99 | 72928 | 70344 | 1.04 |
| 3 | 15129 | 14471 | 1.05 | 30653 | 29684 | 1.03 |
| 4 | 11192 | 10262 | 1.09 | 11909 | 11294 | 1.05 |

| | | | | | | |
|---|---|---|---|---|---|---|
| 5 | 9137 | 7736 | 1.18 | 4351 | 4032 | 1.08 |
| 6 | 6108 | 4847 | 1.26 | 1656 | 1367 | 1.21 |
| 7 | 3706 | 2889 | 1.28 | 635 | 385 | 1.65 |
| 8 | 2153 | 1775 | 1.21 | 434 | 144 | 3.01 |
| 9 | 1564 | 1044 | 1.50 | 253 | 103 | 2.46 |
| 10+ | 8058 | 2197 | 3.67 | 206 | 25 | 8.24 |

**Table S8. Triplex and G-quadruplex motifs are enriched at actual breakpoints**
Although both triplex and G-quadruplex (GQ) motifs are enriched around actual, cancer-associated breakpoints (BPs) compared to control BPs, the difference is more pronounced for triplex motifs on average. Most BPs were not associated with any motif and for both GQ and triplex motifs the results are strongest in the subset of BPs associated with multiple motifs. The pseudo-median and confidence interval (CI) was calculated by Wilcoxon rank sum test in R.

**triplex min score=12**

| **Species** | **Subgroup** | **Triplex (count)** | **Correlation vs MLS** | **Sample (N)** |
|---|---|---|---|---|
| mammals | **low** | 4.2 | **0.217** | 230 |
| mammals | **high** | 6.1 | **-0.444** | 172 |
| birds | **low** | 5.2 | -0.136 | 123 |
| birds | **high** | 9.1 | **-0.239** | 68 |
| fish | **low** | 6.3 | **0.459** | 59 |
| fish | **high** | 9.3 | -0.124 | 78 |

**triplex min score=15**

| **Species** | **Subgroup** | **Triplex (count)** | **Correlation vs MLS** | **Sample (N)** |
|---|---|---|---|---|
| mammals | **low** | 4.2 | 0.046 | 230 |
| mammals | **high** | 6.1 | **-0.280** | 172 |
| birds | **low** | 5.2 | -0.142 | 123 |
| birds | **high** | 9.1 | **-0.384** | 68 |
| fish | **low** | 6.3 | -0.183 | 59 |
| fish | **high** | 9.3 | -0.064 | 78 |

**Table S9. Correlation between triplex motifs and MLS is strongest in species with many triplex motifs in their mtDNA**

Triplex (count) refers to the mean number of mtDNA triplex motifs in the corresponding subgroup. Each “low” subgroup is composed of orders with triplex counts below the median for the whole phylogenetic class, and the “high” group of orders with triplex counts above the median. When we pool data from mammalian orders with fewer triplex motifs than the median for all mammals (n=172 species) we find an inverse relationship with maximum lifespan (MLS) and the same is true for birds. Significant correlations are bolded and underlined (p<0.05).

| Motif | $IR^+$ | $IR^-$ | ratio ($IR^+$ vs $IR^-$) | p-value |
|---|---|---|---|---|
| $ER^+$ (6-15bp) | 18 | 53 | 0.33 | |
| $GQ^+$ (default) | 8 | 106 | 0.07 | p<0.001 |
| $DR^+$ (6-15bp) | 39 | 418 | 0.09 | p<0.001 |

**Table S10. Colocalization of ER, GQ and DR motifs**

Everted repeat (ER) motifs preferably colocalize with inverted repeat (IR) motifs. Neither G-quadruplex (GQ) nor direct repeat (DR) motifs show such an enrichment around IR motifs. Significance based on Fisher’s exact test. Deletion data from the MitoBreak database.
